# Rapid characterisation of hERG channel kinetics I: using an automated high-throughput system

**DOI:** 10.1101/609727

**Authors:** Chon Lok Lei, Michael Clerx, David J. Gavaghan, Liudmila Polonchuk, Gary R. Mirams, Ken Wang

**Affiliations:** Computational Biology, Department of Computer Science, University of Oxford, Oxford, United Kingdom; Pharma Research and Early Development, Innovation Center Basel, F. Hoffmann-La Roche Ltd., Basel, Switzerland; Centre for Mathematical Medicine and Biology, School of Mathematical Sciences, University of Nottingham, Nottingham, United Kingdom

## Abstract

Predicting how pharmaceuticals may affect heart rhythm is a crucial step in drug-development, and requires a deep understanding of a compound’s action on ion channels. *In vitro* hERG-channel current recordings are an important step in evaluating the pro-arrhythmic potential of small molecules, and are now routinely performed using automated high-throughput patch clamp platforms. These machines can execute traditional voltage clamp protocols aimed at specific gating processes, but the array of protocols needed to fully characterise a current is typically too long to be applied in a single cell. Shorter high-information protocols have recently been introduced which have this capability, but they are not typically compatible with high-throughput platforms. We present a new high-information 15 s protocol to characterise hERG (Kv11.1) kinetics, suitable for both manual and high-throughput systems. We demonstrate its use on the Nanion SyncroPatch 384PE, a 384 well automated patch clamp platform, by applying it to CHO cells stably expressing hERG1a. From these recordings we construct 124 cell-specific variants/parameterisations of a hERG model at 25 °C. A further 8 independent protocols are run in each cell, and are used to validate the model predictions. We then combine the experimental recordings using a hierarchical Bayesian model, which we use to quantify the uncertainty in the model parameters, and their variability from cell to cell, which we use to suggest reasons for the variability. This study demonstrates a robust method to measure and quantify uncertainty, and shows that it is possible and practical to use high-throughput systems to capture full hERG channel kinetics quantitatively and rapidly.

**Statement of Significance:** We present a method for high-throughput characterisation of hERG potassium channel kinetics, via fitting a mathematical model to results of over one hundred single cell patch clamp measurements collected simultaneously on an automated voltage clamp platform. The automated patch clamp data are used to parameterise a mathematical ion channel model fully, opening a new era of automated and rapid development of mathematical models from quick and cheap experiments. The method also allows ample data for independent validation of the models and enables us to study experimental variability and propose its origins. In future the method can be applied to characterise changes to hERG currents in different conditions, for instance at different temperatures (see Part II of the study) or under mutations or the action of pharmaceuticals; and should be easily adapted to study many other currents.

## INTRODUCTION

The *human Ether-à-go-go-Related Gene* (*hERG*) is of great interest in the field of cardiac safety pharmacology. hERG encodes the pore-forming alpha subunit of the ion channel Kv11.1 which conducts the rapid delayed rectifier potassium current, *I*_Kr_ (1). Reduction of *I*_Kr_ by pharmaceutical compounds or mutations can prolong the ventricular action potential (2), increase the QT interval on the body-surface electrocardiogram, and is associated with elevated risk of Torsade de Pointes (TdP) (3). Current pharmaceutical regulatory guidelines require the evaluation of effects on the hERG channel as part of pre-clinical drug development (4).

High-throughput automated patch clamp screening for ion current inhibition by pharmaceutical compounds has been widely used to inform pro-arrhythmic safety in early drug discovery. Inhibition data from multiple ion channels can be integrated together using a mechanistically-detailed *in silico* electrophysiology model to predict pro-arrhythmic risk (5). Such a strategy, combining high throughput *in vitro* and *in silico* approaches, is being advocated by a Food and Drug Administration-led initiative, the Comprehensive in vitro Proarrhythmia Assay (CiPA) (6), as a core pillar of future pro-arrhythmic safety assessment. High-throughput automated patch clamp has also been used to characterise the kinetics of a large number of *KCNQ1* mutants that were previously variants of unknown significance (7).

Mathematical modelling of ion channel kinetics provides a quantitative summary of our current understanding, and can serve as a powerful predictive tool. The parameters in ion current models can be biophysically and physiologically meaningful, and are therefore of interest in their own right. Parameterisation (or *calibration*) of mathematical models is a concise way to characterise ion current kinetics, and can also be used to quantify variability between experiments (8). A wide range of models have been proposed to describe *I*_Kr_, with varying levels of biophysical detail and numbers of parameters (see Beattie et al. (9, Appendix A)). Until we have a full and clear understanding of the underlying mechanisms, simple models that capture the most relevant characteristics with a small number of parameters may be preferred.

Voltage clamp experiments are a common source of data for calibrating ion channel models. The first models of ionic currents were proposed by Hodgkin and Huxley (10), who used step-wise voltage protocols to isolate and measure different aspects of ionic currents (e.g. time constants and voltage-dependent steady states). Following in their footsteps, many voltage step protocols have been designed to highlight particular current kinetics. Typically, these protocols involve long sections during which the channels are brought into a particular state, before a brief interval during which a current is measured and then summarised using either a peak current or by fitting an exponential curve and deriving a time constant. By design, these protocols focus on a single aspect of an ion current so several such protocols are needed to parameterise a model fully. More recently, simulation experiments have shown that condensed voltage clamp protocols can be used to provide the required information in a much shorter time (11, 12). A study by Beattie et al. (9) demonstrated *in vitro* that sinusoidal protocols can be used to rapidly (8 seconds) characterise hERG kinetics on a manual patch clamp setup. Due to hardware limitations, some automated high-throughput systems can only perform square wave or ramp voltage clamp protocols. Here we extend the approach of Beattie et al. (9) to make it applicable to such automated high-throughput patch clamp systems.

Efforts have been made to address the variability observed in measurements of the hERG channel (13). However, the variability of baseline hERG characteristics remains incompletely understood. Understanding and quantifying this variability, whether it be due to cell-to-cell variability (also known as *extrinsic variability* or *population variability*) or to observational errors/uncertainties, is crucial in establishing the credibility and applicability of model predictions (14). Quantifying the variability in hERG channel kinetics requires a large number of high-quality patch clamp measurements and an appropriate statistical framework. The duration of a standard combination of protocols makes it difficult to use them to fully characterise the current in a single cell, so that reaching the required number of cells for a thorough statistical analysis would be a very difficult and time consuming task.

We present a new approach to overcome this problem by using a novel protocol and a high-throughput system to rapidly record many cells’ kinetics in parallel. Using these methods, we construct 124 cell-specific parameterisations of a hERG model, and validate all of our model predictions against a set of independent protocols that have not been used in training/fitting the model. To ensure the stability and reproducibility of our results *within the same cells*, we repeat all of our measurements twice. We employ a hierarchical Bayesian framework (a multi-level statistical modelling technique) to describe the variability of hERG channel conductance and kinetics between cells, and to infer the covariance between the model parameters across different cells. This study greatly increases the utility of automated high-throughput systems, and provides robust tools for the uncertainty quantification that comprise an essential component of an *in silico* assay.

## MATERIALS AND METHODS

We began our work with a synthetic data study to inform the experimental design of the voltage protocols, and applied inference techniques to assess the amount of information such protocols can provide. The motivation and rationale of our newly designed protocol are discussed in the Experimental methods section. Experiments using this new protocol were performed on the Nanion SyncroPatch 384PE platform (Nanion Technologies GmbH, Germany) with a temperature control unit. We then applied global optimisation, Markov Chain Monte Carlo (MCMC), and hierarchical Bayesian techniques to recover parameters for a mathematical ion current model for each individual cell, as described below.

### Mathematical model

We used a recently published hERG model by Beattie et al. (9), which has a Hodgkin & Huxley-style structure. This model structure has been widely used in many studies with slight modifications: the root of the model traces back to Zeng et al. (15), where the same model structure is used but with the inactivation gate modelled as an instantaneous steady-state response. Later in the Ten Tusscher et al. (16) model, the same model structure was used, but extra parameters were introduced to make the time constant independent of the steady state. In the model that we use, the current, *I*_Kr_, is modelled with a standard Ohmic expression,

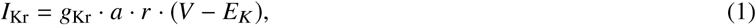

where g_Kr_ is the maximal conductance, *a* is a Hodgkin and Huxley (10) activation gate, and *r* is an inactivation gate. *E*_*K*_ is the reversal potential, also known as the Nernst potential. *E*_*K*_ was not inferred but was calculated directly using

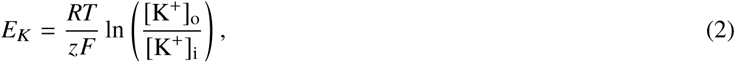

where *R* is the ideal gas constant, *T* is the absolute temperature, *F* is the Faraday constant, and *z* is the valency of the ions (equal to 1 for K+). [K+]_o_ and [K+]_i_ denote the extracellular and intracellular concentrations of K^+^ respectively, which were determined by the experimental solutions as 4 mM and 110 mM respectively. The model structure is shown in Figure 1, where

**Figure 1:**
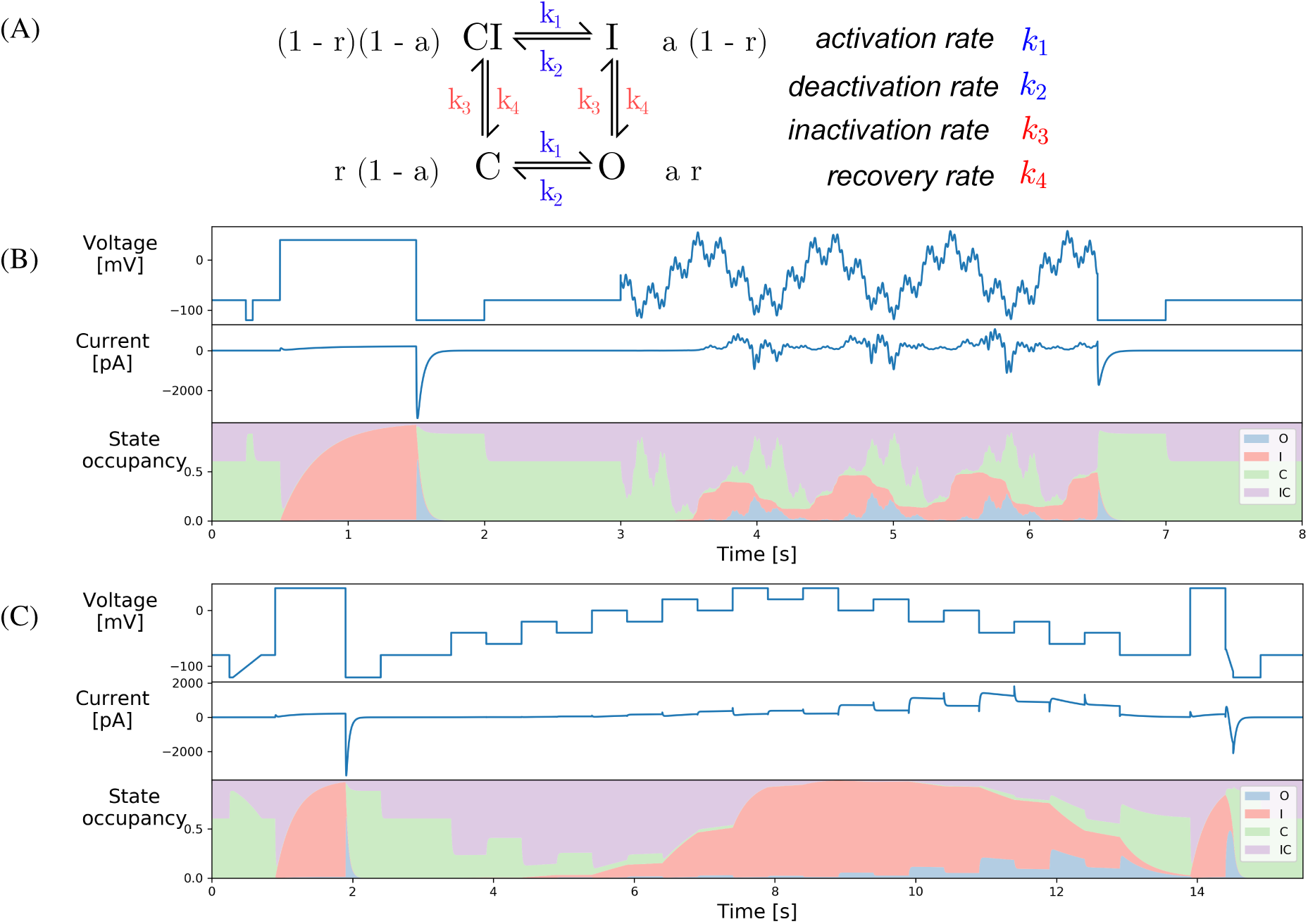
**(A)** The Hodgkin-Huxley model structure shown in equivalent Markov state diagram format. Four states are linked with arrows with rate transitions given by the expressions on the right. The probabilities of each state are given next to them in terms of the Hodgkin-Huxley gates *a* and *r*. **(B)** A manual patch clamp protocol comprised of an 8 second voltage clamp protocol designed for rapid characterisation of ion channel kinetics by Beattie et al. (9). **(C)** Our novel 15-second protocol, which we term the ‘*staircase protocol*’, designed for any patch clamp set-up including high-throughput automated systems, that is similarly able to characterise the full kinetics of the hERG channel model. Both (B) and (C) show the voltage protocol (top panel), an example of the simulated current using the room temperature parameters from Beattie et al. (9, cell #5) (middle panel), and the corresponding state occupancy (bottom panel).

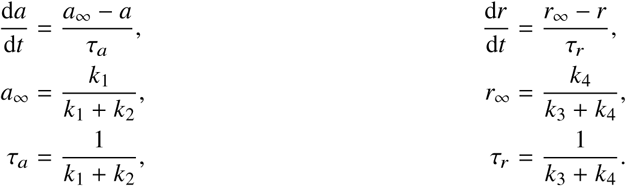

Where

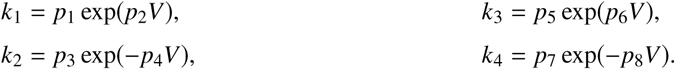

Our model consists of 9 positive parameters ***θ*** = {*g*_Kr_, *p*_1_, …, *p*_8_}, each of which must be inferred from the experimental data.

Simulations were run using Myokit (17), with tolerance settings for the CVODE solver (18) set to `abs_tol`= 10^−8^ and `rel_tol` = 10^−10^. All codes and data are freely available at https://github.com/CardiacModelling/hERGRapidCharacterisation.

### Statistical model and parameter inference

To infer model parameters from experimentally observed data under a probabilistic and Bayesian framework, we specified a statistical model to relate the mathematical model and the observed experimental data:

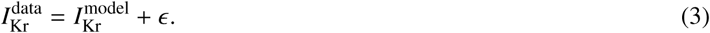

We assumed that noise arises from a normal distribution *ϵ* ∼ 𝒩(0, *σ*^2^). This is equivalent to writing 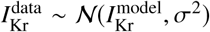, which allows us to formulate the likelihood of observing the data **y** given parameters ***ϕ*** = ln(***θ***) as

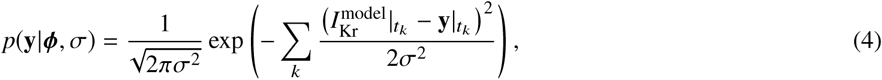

where *t*_*k*_ is the time point at index *k*. We chose the parameter transformation ***ϕ*** = ln(***θ***) to turn our positively constrained physical model parameters to be unconstrained optimisation variables. Using Bayes’ theorem, we can now write an equation for the likelihood of a parameter set given the observed data (the posterior) as

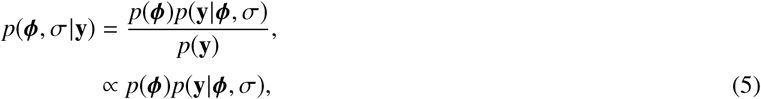

with the prior

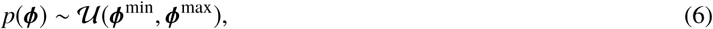

where **𝒰** (·) represents a uniform distribution.

Here **y** was assumed to be the 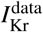 in Eq. 3, after leak correction and E-4031 subtraction have been applied as part of the data processing. We chose a uniform prior, and expected our posterior to be dominated by the observed data. The details of the choice of ***ϕ***^min^, ***ϕ***^max^ are given in Section S6.2.2 in the Supporting Material. Such a formulation extends our model parameters to {***θ***, *σ*^2^}, to fully describe both the biophysical and statistical models.

We used a two-step approach to infer the model parameters. Firstly, we used a global optimisation algorithm (19) to identify the parameters. Secondly, we utilised a Monte-Carlo based sampling scheme to obtain the posterior distribution, using a population MCMC (20) algorithm with adaptive Metropolis (21) as the base sampler. The benefits of this approach are twofold. First, using a Bayesian framework allows us to incorporate prior knowledge. Second, we construct a probability (posterior) distribution to quantify uncertainty in the parameter set due to noise in the data. All inference and sampling were done via our open source Python package, PINTS (22).

### Hierarchical Bayesian model

We combined multiple experimental recordings using a multi-level modelling technique known as a hierarchical Bayesian model. Under this framework, we assume the vector of the transformed parameters ***ϕ*** for a particular cell follows a multivariate normal distribution which describes how these parameters are distributed between all cells, namely ***ϕ*** ∼ 𝒩(***µ***, **Σ**). Given our choice of parameter transformation, this is equivalent to writing ***θ*** ∼ LogNormal(***µ***, **Σ**), that is the vector of parameters ***θ*** for a particular cell follows a multivariate log-normal distribution. Then we used the hierarchical Bayesian model to infer the mean vector ***µ*** and covariance matrix **Σ** across cells, and hence determined any correlation in model parameter sets between cells. The parameter dependency for this hierarchical Bayesian model is shown in Figure S5 in the Supporting Material.

The full hierarchical Bayesian likelihood ℒ was specified as the product of: (a) the probability of producing data **y**_*j*_ on each cell given the parameter vector for each cell ***θ*** _*j*_ and noise *σ*_*j*_; (b) the probability of obtaining each individual well parameter set ***θ*** _*j*_ from the ‘top-level’ LogNormal distribution across wells defined by the hyperparameters; and (c) the priors — the prior of the hyperparameters (also known as the ‘hyper-prior’) and the prior of *σ*_*j*_. That is,

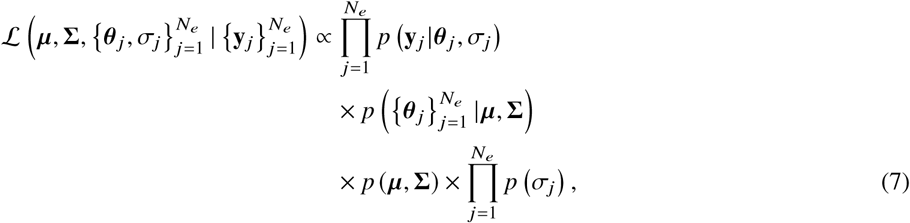

where ***µ***, **Σ** are the hyperparameters of the hierarchical model representing the mean vector and covariance matrix of the individual ‘low-level’ parameters, and 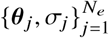 are the set of individual ‘low-level’ parameters for each of the *N*_*e*_ repeats of the experimental recordings 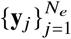.

We sampled the full hierarchical Bayesian model using a simplified version of the Metropolis within Gibbs (MwG) (23) method which we have termed ‘*pseudo-MwG*’ (see Section S6 and Figures S9 in the Supporting Material, but note this simplification is only applicable for our particular setting — where the number of data points in the time traces vastly outweighs the number of cells). We also describe the details of the choice of likelihoods and priors and sampling algorithms in Supporting Material Section S6, and we test the LogNormal distribution assumption in Section S8.

We used the inferred covariance matrix ****Σ**** to study the correlation (corr(***θ***)) between the model parameters, which are related by

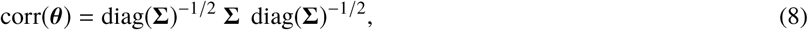

where diag(·)^−1/2^ denotes the square root of the matrix of the diagonal entries. The posterior predictive distribution *p*(***θ*** |…) allows us to make predictions about how future experiments will behave, where (…) indicates all other variables appearing in Eq. 7. It can be computed using

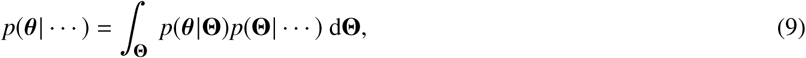

where **Θ** = {***µ***, **Σ**}, a concatenation of all the individual hyperparameters within ***µ*** and **Σ**. The integration was approximated by summing over the probability density functions which are defined by the samples of **Θ**.

### Synthetic data studies

Before implementing experiments, we confirmed the identifiability of model parameters using our protocols and parameter inference algorithms through a synthetic data study. We generated synthetic data (with added synthetic noise) with some known ‘true’ parameters ***θ***^true^. First, we used the synthetic data to design and optimise our protocols and to ensure that the protocols give access to sufficient information for parameter characterisation. Second, we assessed our inference methods, described in the previous section, by asking how confident we are in our inferred parameters. In Supporting Material Section S6.3.1, we show that our newly designed protocol, the *staircase protocol* (see Figure 1C), is information-rich, in that we are able to fully recover the ‘true’ parameter in a synthetic data study using our protocol.

We also tested our hierarchical Bayesian model, to ensure that it is possible to infer the underlying distribution of the parameters. We generated our individual synthetic data from a predefined multivariate normal distribution, where parameters are correlated. In Supporting Material Section S6.3.2, we applied our hierarchical Bayesian model analysis to the synthetic data, assuming we did not know the underlying covariance between parameters, and we were able to reconstruct the correlation matrix of our predefined distribution with very high accuracy. This provides us with confidence that our method is able to correctly infer the underlying correlation between parameters. We describe the rationale and procedure of the synthetic data study in detail in the Supporting Material Section S6.

### Experimental methods

Whole-cell patch-clamp voltage clamp experiments were performed on Chinese Hamster Ovary (CHO) cells stably transfected with hERG1a (Kv11.1), with temperature control set to 25 °C, using the Nanion SyncroPatch 384PE platform (Nanion Technologies GmbH, Germany). The temperature of the system’s ‘cell hotel’ was set to ∼15 °C. The machine is an automated high-throughput platform, in which each run (or chip) is able to measure up to 384 wells (with one cell per well) simultaneously. Single hole chips with medium resistance (Nanion order number #221102) were used. Solutions used in all measurements are provided in Table S2 in the Supporting Material.

A schematic of the experimental procedure is shown in Figure 2, which shows the voltage clamp protocols used in the experiments. A total of of 9 voltage clamp protocols were used, including (green) our newly developed staircase protocol, (blue) an activation current-voltage (I-V) protocol, a steady-state inactivation I-V protocol, a hERG screening protocol, a delayed afterdepolarization (DAD)-like protocol, an early afterdepolarization (EAD)-like protocol, and action potential-like protocols with beating frequency 0.5 Hz, 1 Hz and 2 Hz, as shown in Figure 2. Note that, due to the automated platform, the action potential-like protocols have to be comprised of a series of linear ramps and steps rather than curves. Details of the protocols are given in Section S1 in the Supporting Material. Every protocol (the entire procedure in Figure 2) was applied to every well.

**Figure 2:**
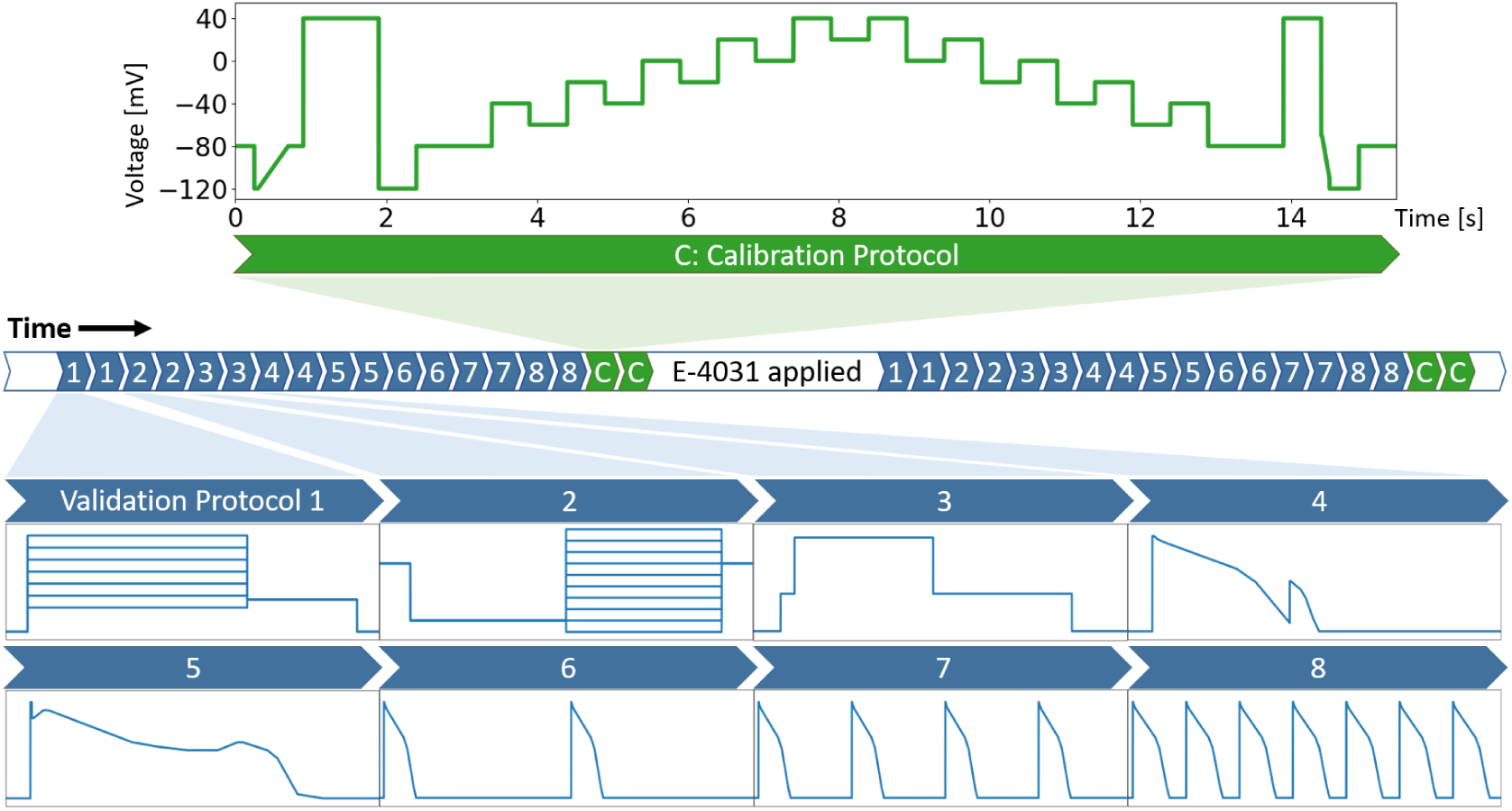
A schematic of the experimental procedure showing the sequence of voltage clamp protocols used. A total of 9 voltage clamp protocols were used, each of them was performed 4 times: twice before E-4031 addition, and twice after, to ensure stability and reliability of the recordings. Only the staircase protocol (green, 15 seconds) was used for fitting (or calibrating) the mathematical model. All of the other 8 protocols (blue) were used for validation only. White sections indicate a non-measurement region, where cells were held at −80 mV, to allow the cells to settle to steady state between protocols (>5 s), or to allow the drug to wash in (>5 min). For details of the protocols please refer to Supporting Material (Section S1).

Only the staircase protocol (green) was used in fitting (or calibrating) the mathematical model. We show that we can fully characterise *I*_Kr_ for each cell using just this one protocol, as our staircase protocol is information-rich. A comparison between the staircase protocol and a previously developed protocol (9) is shown in Figure 1B and C. However, due to hardware limitations, the previous protocol does not work in most high-throughput automated systems as they cannot perform clamps to arbitrary time-varying functions and are limited to ramps and steps. Hence a similar idea from Beattie et al. (9) — using an information-rich protocol — was adapted, and the rationale of our staircase protocol is discussed below. We designed the staircase protocol with only voltage steps and ramps, such that it is applicable to any patch clamp machine including the high-throughput automated systems.

A demonstration that a mathematical model is able to reproduce the experimental training data is not sufficient to conclude that it is a good representation of ion channel kinetics — in particular, we may be uncertain how well the model performs under physiological conditions. The fitted models for each cell were therefore validated by comparing with experimental data from each of the other 8 protocols (blue in Figure 2). Our validation set consists of 1) two traditional I-V protocols together with a simple hERG activation step and 2) five physiologically-inspired protocols that mimic cardiac action potentials. The first set allows us to compare with the traditional approach. More importantly, the second set allows us to have confidence in predictions of *I*_Kr_ responses, which is particularly useful when an ion channel model is embedded in a cardiac action potential model. This series of validations allows us to demonstrate that the models fitted using this new protocol yield trustworthy cell-specific predictions.

### Protocol design

The underlying rationale of the *staircase protocol* is to force the protocol to explore the full dynamics of the system at different voltage values, over a physiologically relevant voltage range. By observing the changes in the current after each step, the voltage dependency of the channel at that particular voltage can be deduced. Each voltage step is held for 500 ms, which is chosen to be long enough to observe the characteristic decay of *I*_Kr_. Therefore, by going through different step-ups and -downs, the protocol explores the dynamics at different voltage values, and hence our statistical inference method is able to infer the underlying model parameters.

Two ramps are implemented before and after the main staircase. The first one is used to estimate the leak current, see the next section for more details. The second one is designed to estimate experimentally the reversal potential *E*_*K*_ by having a ramp that quickly crosses the expected *E*_*K*_, which we expect to be in the range of −70 mV to −100 mV. However, the hERG channel closes relatively quickly at low voltage, during which only a very small *I*_Kr_ signal can be recorded. We therefore implemented a large step up to 40 mV before the ramp to open the channel, so that we can record a high signal-to-noise ratio *I*_Kr_ trace that goes from positive to negative. An example demonstrating the use of the two ramp design is shown in Figure S2 in the Supporting Material. Implementation of all of the above leads to the staircase protocol shown in Figure 2.

### Post-processing experimental data

We assumed that our observed current from hERG CHO cells under control conditions is

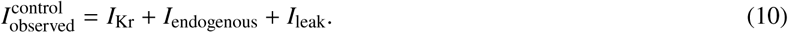

To ensure the currents recorded are purely *I*_Kr_, we performed the series of corrections described below. First, leak corrections were applied to all measurements to eliminate *I*_leak_. Second, E-4031 subtraction was applied to remove *I*_endogenous_ (the sum of any native voltage-dependent ion currents that were present in CHO cells besides the overexpressed hERG). We also describe our partially automated quality control criteria to select and ensure high quality *I*_Kr_ recordings.

### E-4031 subtraction

To eliminate any endogenous voltage-dependent background currents within the hERG CHO cells (*I*_endogenous_ in Eq 10) we measured the full set of 9 voltage protocols twice, see Figure 2; once with Dimethyl sulfoxide (DMSO) vehicle conditions in which 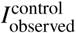 in Eq. 10 was measured, and once under the addition of 0.5 µM E-4031, a hERG channel selective blocker with IC_50_ value ≲10 nM, so that

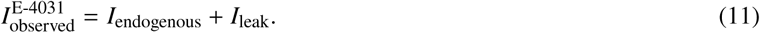

As shown in Figure 2, a period of ∼5 min was allowed for the E-4031 block to reach equilibrium. All currents shown or used in this study are the leak corrected currents measured in control conditions minus the leak corrected currents that remained after E-4031 addition, which we assume yields uncontaminated *I*_Kr_.

### Leak correction

A leak step was implemented to infer the leak current, where we assumed

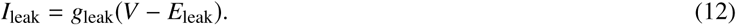

Here, g_leak_ is the leak current conductance and *E*_leak_ is the leak current reversal potential. Since this model is linear we can estimate its parameters g_leak_, *E*_leak_ in two ways; either by using two points, or better by using a linear ramp. Then our final *I*_Kr_ is given by

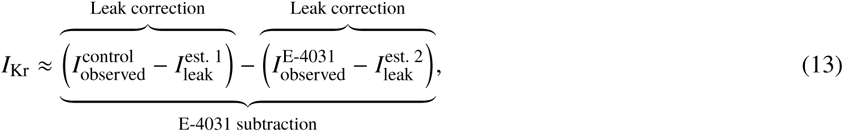

where 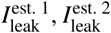 are leak currents estimated using Eq. 12.

We assumed that at −80 mV, *I*_Kr_ is fully closed and will not be opened by going to a voltage below −80 mV. We therefore implemented a linear ramp from −120 mV to −80 mV, as seen in the first 1 second of the staircase protocol (green), in Figure 2. All non-zero current measured during the ramp was assumed to be leak current in the form of Eq. 12, and a linear regression was used to fit its I-V relation and to obtain the leak model parameters. We show in detail the use of our linear ramp to infer the leak model parameters in Figure S2, in which we can see that the recorded current during the ramp shows a reliably good linear relation. Therefore, this leak ramp can be used to check the linearity of the leak current in our model, that is the linearity in its I-V relation, which cannot be achieved using the standard two-point method for leak estimation.

In a similar fashion, for *all* validation protocols, instead of a linear ramp, a traditional two-point method was used. This method was implemented and performed automatically by the platform we used. A leak-step from −80 mV to −100 mV was used to leak-correct the experimental data. Such a leak-step was performed before every measurement to correct the recording that followed. However, we noticed that some of these leak corrections can ‘over-correct’ the current. That is, since we assumed *I*_Kr_ is only negative when the voltage is below its reversal potential, which was about −85.2 mV, if the leak-corrected current showed a negative current at voltages substantially larger than the reversal potential, then we would expect the automated system to have overestimated the leak current. Such over-correction was most noticeable during the highest voltage step during the protocol, where *I*_leak_ was at its maximum and *I*_Kr_ was usually almost zero. When this occurred, we re-estimated the leak correction by adding an extra linear leak current of the form 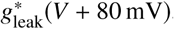, where 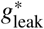 was chosen such that the leak-corrected current during the highest voltage point was zero *if negative before*. Due to the linearity, the final leak correction remains equivalent to Eq. 12 with different parameters.

### Partially automated quality control

After the experiments, we applied a strict set of criteria as an automated selection process for quality control of our experimental data. The details of our criteria are summarised in Table 1. We applied a strict cut-off for seal resistance (R_seal_), cell capacitance (C_m_), and series resistance (R_series_) through the whole set of measurements, set by our first quality control criterion (QC1). QC2 required a high signal-to-noise ratio (SNR) recording, such that our measurements contained enough useful information for model inference. We also compared the stability of the recordings in QC3, where each protocol consisted of two measurements recorded in the same cell that must be similar and stable. QC4 required R_seal_, C_m_, and R_series_ to be stable before and after E-4031 addition. QC5 required that the addition of hERG blocker E-4031 must reduce a certain amount of the recorded current, to ensure that our recordings consisted mainly of *I*_Kr_ even before adding the blocker. Finally, over-correction of leak can occur during high voltage steps, as discussed in the previous section, QC6 ensured that no negative current occurred at voltages substantially larger than the reversal potential. Note that, QC1 to QC4 are general criteria that are advocated to be used in all whole cell patch-clamp voltage-clamp experiments, whereas QC5 and QC6 contain prior knowledge of *I*_Kr_ and are tailored to hERG measurements.

**Table 1:**
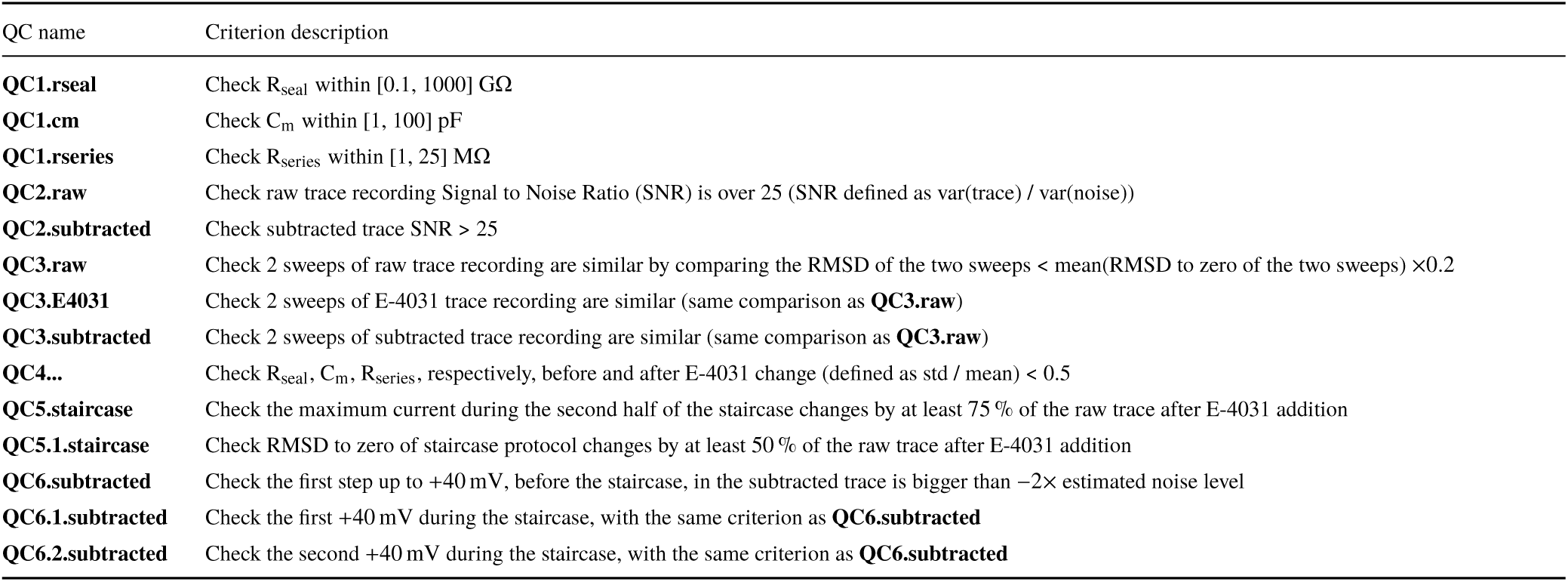
A summary of the fully automated quality control criteria for the staircase protocol, QC1–QC6. RMSD = Root Mean Square Difference.

Using our automated high-throughput system, we recorded a total of 384 well recordings. Our automated quality control removed 173 wells, leaving 211 well recordings. We then manually checked all the recordings, and subsequently removed a further 28 wells that did not look anything like the rest of the 183 cells, 6 typical examples are shown in Figure S4 in the Supporting Material. Therefore our automated quality control has achieved >86 % positive predictive value. The machine’s ‘standard’ quality control selects wells mainly based on the R_seal_, C_m_, and R_series_ values, which were set to the same values we used in our automated QC1 in Table 1, and removed only 46 wells (which were all within our 173 discarded wells). Further comparisons and details of our automated quality control results are shown in Supporting Material Section S5. Our automated quality control is available at our GitHub repository.

Our mostly-automated quality control was applied only to the staircase calibration protocol. In this study, we further require our validation data to contain high quality validation recordings. We therefore manually selected 124 cells within our 183 cells that passed our quality control and hence have good recordings for both calibration and validation protocols; this ensures the quality of the experimental data used in this study. With that, we conclude our success rate of recording our staircase protocol is 183 out of 384 wells and for the full dataset is 124 out of 384 wells, which can be performed within one hour.

## RESULTS

### High-throughput experimental recordings

Figure 3 shows the voltage clamp recordings measured with the 9 different protocols, and the corresponding voltage protocols. All results shown are the first of the two repeats of our recordings. Our analysis was repeated for the second of the two repeats to ensure the reproducibility of our results in the same cells: the intrinsic (within-cell) variability is sufficiently small to be negligible, see Figure S12 in the Supporting Material.

**Figure 3:**
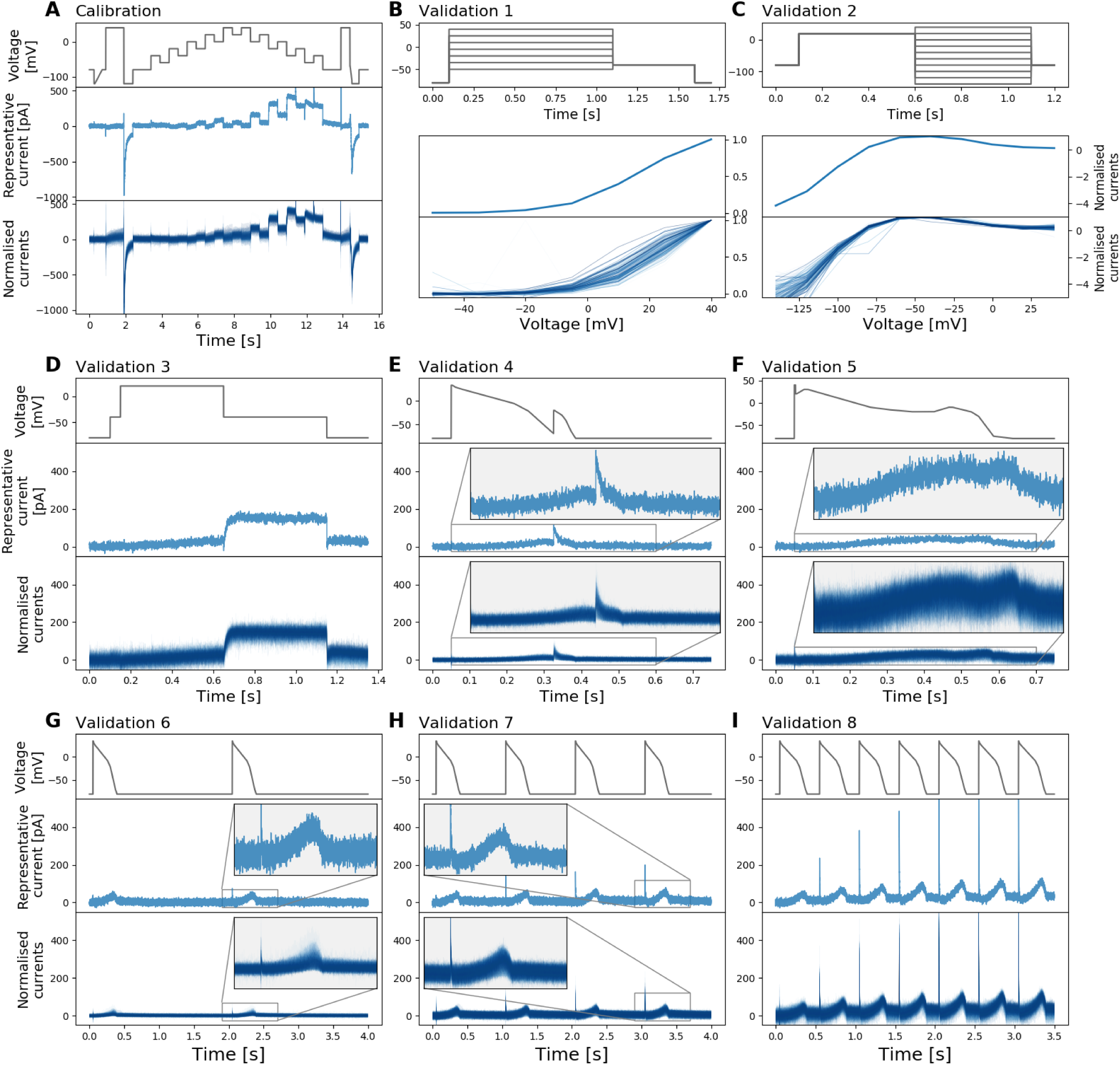
Whole-cell patch-clamp voltage clamp recordings under 9 different protocols which were all measured in each cell. **(A)** Shows the staircase protocol (top panel) in black and the corresponding recording on a single cell (middle panel) and normalised recordings from all 124/384 wells that passed quality control (bottom panel) in blue. Conductance normalisation was done by multiplying each current by a scaling factor to minimise the absolute difference between each trace and a reference trace (middle panel). **(B–I)** The eight different protocols used as validation of the model calibration, which are the activation current-voltage (I-V) protocol, the steady-state inactivation I-V protocol, the hERG screening protocol, the DAD-like protocol, the EAD-like protocol, and the cardiac action potential-like protocol at 0.5 Hz, 1 Hz and 2 Hz, respectively. All experimental recordings, both the single cell (middle) and 124 cells (bottom), are shown in blue, which were measured under the protocol (black) shown in the panels immediately above. In **(B, C)**, validation 1 and 2 show the I-V relations extracted from the currents.

Figure 3A shows the staircase protocol (black) and the corresponding experimental recordings (blue). The middle panel shows the raw current recording of a single cell; the bottom panel shows the normalised current recordings from all 124 wells that passed quality control. Normalisation is applied for visual comparison only, as each hERG-transfected CHO cell is expected to have a different total conductance hence giving a different magnitude of the current recorded. Currents are normalised by scaling them to minimise the absolute difference between each trace and a reference trace (middle panel). Our recordings show a very similar result to the *I*_Kr_ simulation shown in Figure 1, and the simulated result used parameters calculated/fitted completely independently by Beattie et al. (9).

Figure 3B–I show the recordings of the other 8 validation protocols from the *same cells*. The activation step in Figure 3D recorded a typical *I*_Kr_ response, where the step down of voltage to −40 mV largely opens the channels. Figure 3G–I also shows typical *I*_Kr_ responses to the action-potential clamp at different pacing frequencies, where *I*_Kr_ is active during repolarisation of the action potential. Also note the sharp opening of *I*_Kr_ at the upstroke which changes with pacing frequencies and increases dramatically but very consistently across all the recorded cells.

### Individual cell fitting and validation

Figure 4 shows the same voltage clamp recordings (blue) in Figure 3, measured under the 9 different protocols (black), together with model fitting and validation results. All recordings shown were performed on a *single cell*. The mathematical model, shown as red lines, is fitted *only* to the data recorded under the staircase protocol that is shown in Figure 4A. The result of the fitting for a single cell is shown in the middle panel of Figure 4A, demonstrating an excellent fit between experimental measurement and simulated current. The inferred parameters are shown and studied in detail in the next three sections.

**Figure 4:**
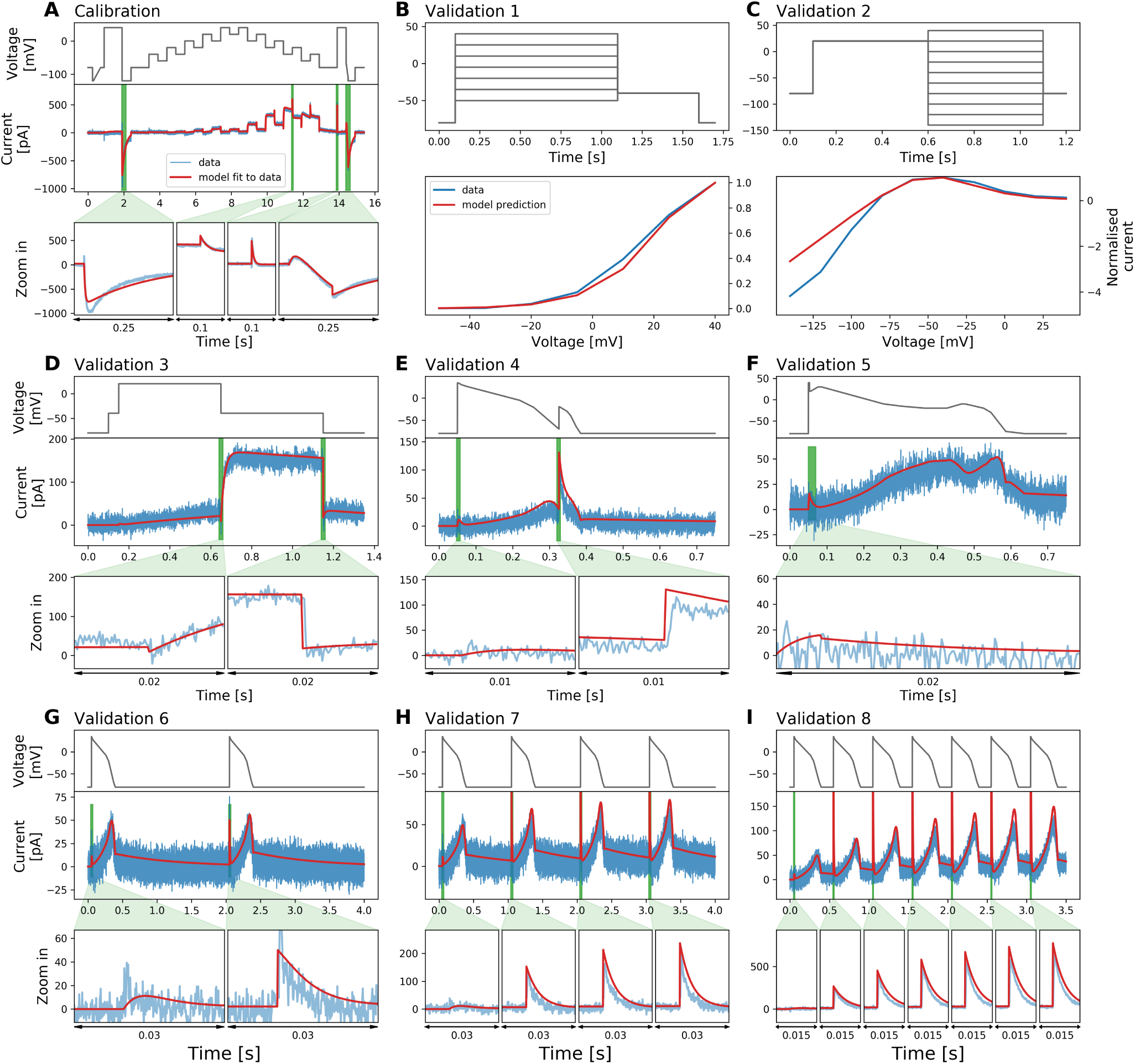
Whole-cell patch-clamp voltage clamp recordings under 9 different protocols which were measured on a single cell, and the model fitting and validation results. **(A)** Shows the staircase protocol (black) and the corresponding recording (blue). The mathematical model is calibrated using this recorded data, and shown as a red line. **(B–I)** The eight different protocols used as validation of the calibrated model, which are the activation current-voltage (I-V) protocol, the steady-state inactivation I-V protocol, the hERG screening protocol, the DAD-like protocol, the EAD-like protocol, and the cardiac action potential-like protocol at 0.5 Hz, 1 Hz and 2 Hz, respectively. All experimental recordings are shown in blue, which were measured under the protocol (black) shown in the panels immediately above, and the validation predictions of the model are shown in red. Zoomed-in image of the green shaded regions are shown underneath each panel to reveal the details of the spikes, in which our model also shows excellent predictions of the faster timescale behaviour. In **(B, C)**, validation 1 and 2 show I-V relations extracted from these protocols.

In Figure 4B–I, we show the results of the validation predictions under 8 other protocols. We validated our trained model by testing its ability to predict independent experimental outcomes under different protocols, which were measured *in the same cell*. All validation predictions were performed by using the inferred parameters in the fit to the staircase protocol (Figure 4A) to simulate the other 8 protocols (Figure 4B–I). The predictions of all the protocols match very well to the experimental data, with the simulated currents giving a close match to the experimental recordings.

The physiologically-inspired voltage clamp protocols (Figure 4E–I) mimic the membrane voltage of the cardiac action potential at normal conditions at different beating rate and EAD/DAD-like conditions. The ability to predict the current response under these physiologically-inspired voltage clamp protocols is particularly important for use in physiological or pharmacological studies. This shows the reliability of the hERG ion-channel model predictions at different physiological conditions, for example, when it is embedded in a whole-cell cardiac model for further predictions.

In Figure 5 we present our model fitting and validation results for all 124 cells, compared against the experimental recordings measured under the 9 different protocols. We applied the same fitting and validation procedure, as used for the single cell discussed above, to all 124 cell measurements. To visualise the variability in only hERG kinetics (and not maximum conductance), we plotted all currents normalised as described in the previous section.

**Figure 5:**
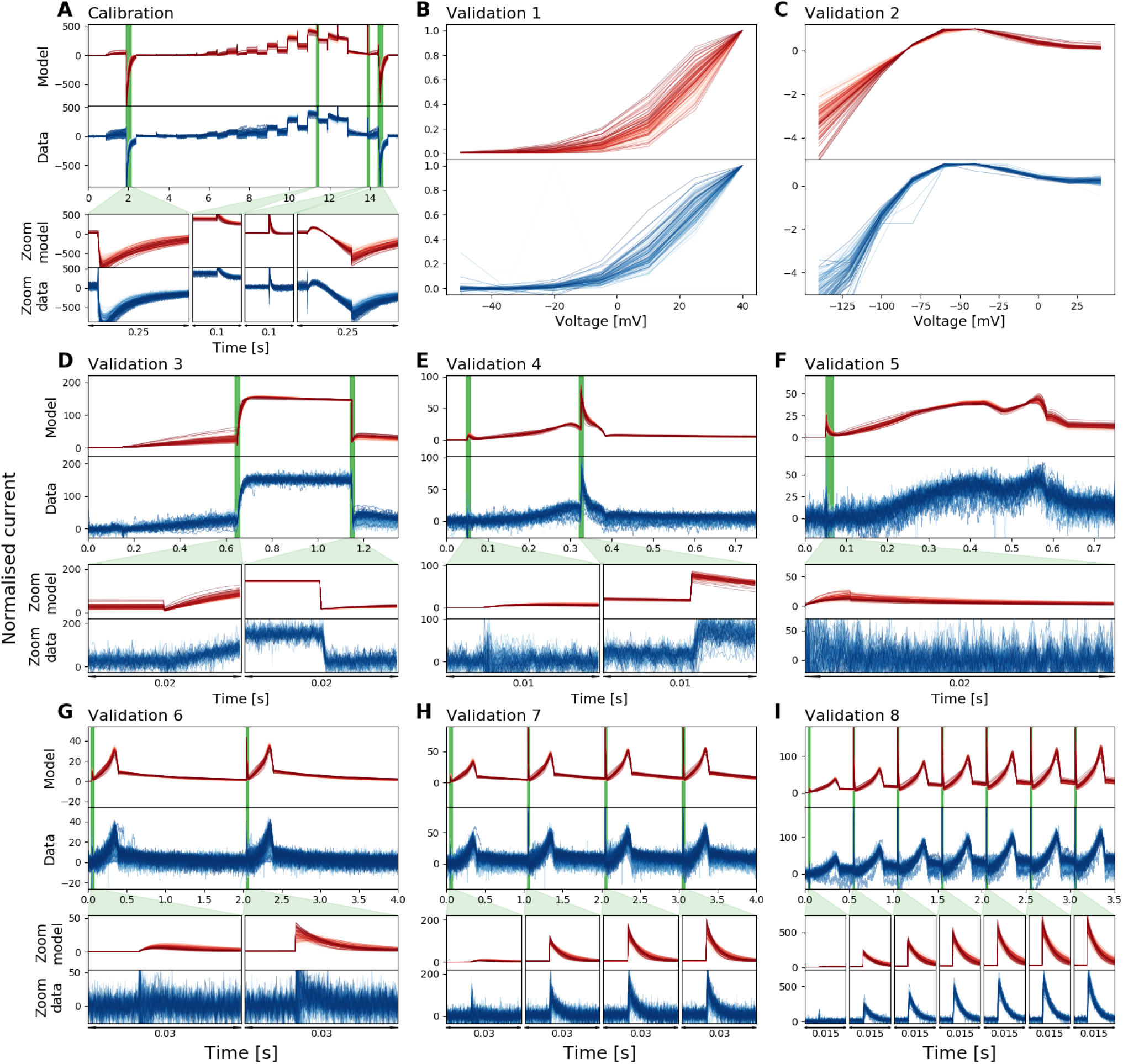
Normalised whole-cell patch-clamp voltage clamp recordings for 124 cells under 9 different protocols, and the model fitting and validation results. All currents are normalised by scaling them to minimise the absolute difference between each trace and a reference trace. From **(A)** to **(I)**: The staircase protocol which is used as the calibration protocol, the activation current-voltage (I-V) protocol, the steady-state inactivation I-V protocol, the hERG screening protocol, the DAD-like protocol, the EAD-like protocol, and the cardiac action potential-like protocol at 0.5 Hz, 1 Hz and 2 Hz, respectively. All the model calibration results and validation predictions are shown in the top panels (red), and are compared against the experimental recordings shown in the bottom panels (blue). Magnifications of the green shaded regions are shown underneath each panel to reveal the details of the spikes, in which our models show extraordinarily good predictions to the details. The normalised current for all protocols are shown except for the activation I-V protocol and the steady-state inactivation I-V protocol where the summary statistic I-V relationships are shown.

We quantified the fits and predictions using relative root mean square error (RRMSE) defined as the root mean square error between the model simulation and the experimental data, divided by the root mean square distance of the data to a zero current trace:

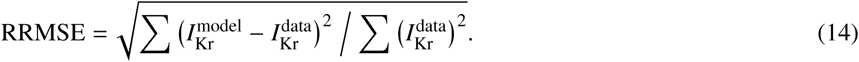

Using this RRMSE quantification, the difference in the absolute size of the current across cells due to varying conductance is eliminated, and RRMSE scores are comparable between cells. Figure 6 shows the RRMSE histograms for all cells and for six of the protocols. Markers indicate the best (∗), median (‡) and 90^th^ percentile (#) RRMSE values, and corresponding raw traces are shown in the three panels above. The marker ♦ indicates the reference cell shown in Figures 3 & 4. There are some small discrepancies in the predictions, for example in Figure 6B in the 90th percentile predictions. But overall these results demonstrate that all our 124 models make very good predictions for the recorded current kinetics.

**Figure 6:**
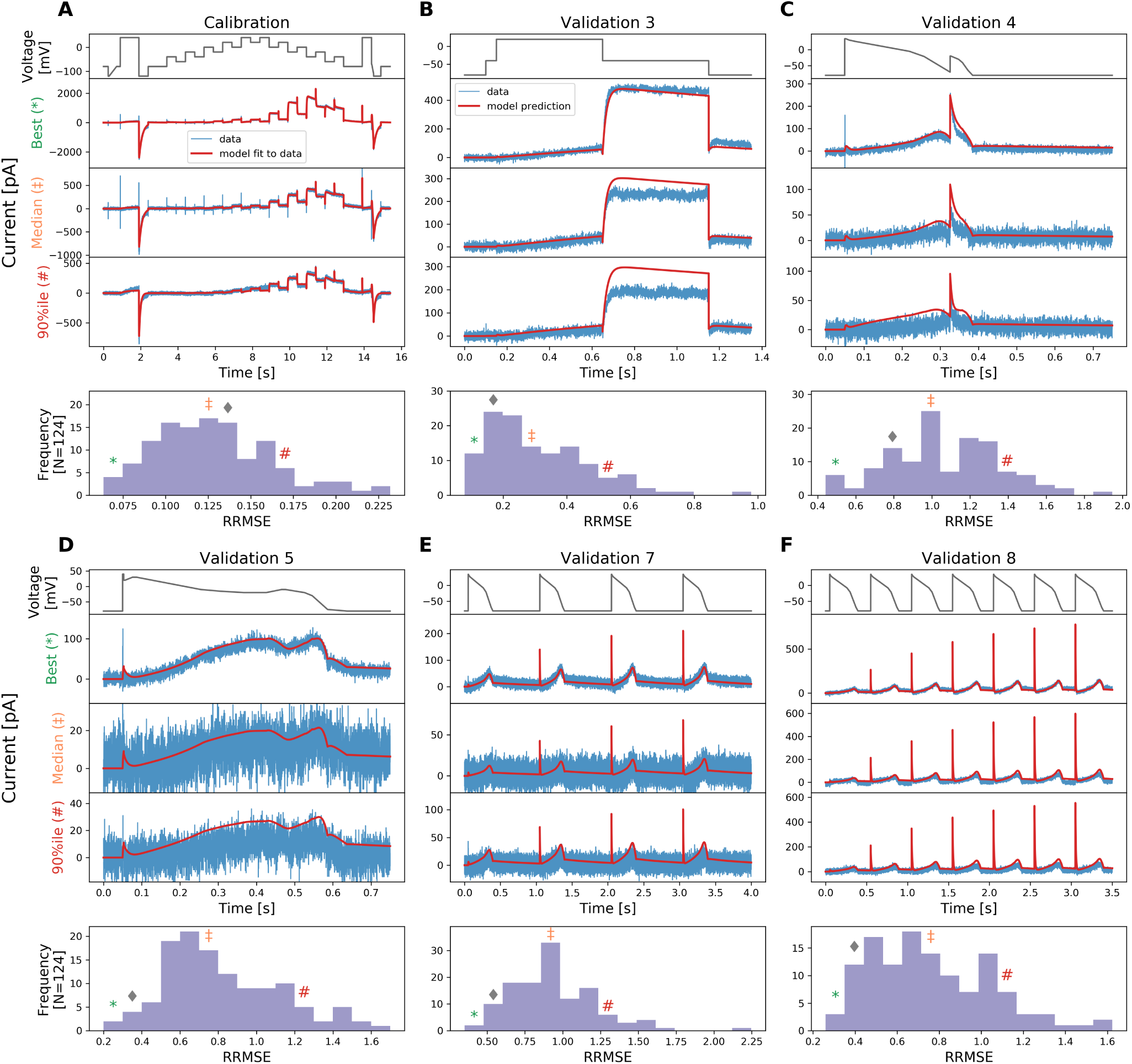
The relative root mean square error (RRMSE, given by Eq. 14) histograms for all 124 cells and for all 6 protocols used. Markers indicate the best (∗), median (‡) and 90^th^ percentile (#) RRMSE values, and marker ♦ indicates the error for the reference traces shown in Figures 3 and 4. For each protocol, the raw traces with the best, median and 90^th^ percentile RRMSE values, for both the model (red) and data (blue) are shown, with the voltage clamp above. Note that the currents are shown on different scales, to reveal the details of the traces.

Next, we first qualitatively inspect the variability in the hERG kinetics measurements. Since we measured the *I*_Kr_ using exactly the same experimental set-up for each cell, we can clearly see the variability *between measurements* in all of the recordings, as illustrated in Figure 5. Different protocols demonstrate different levels of variation. It is clear that, amongst the 6 protocols, the staircase protocol and the two I-V protocols show the strongest variation between measurements.

To investigate this further, we have used our mathematical model to study the variability in the parameter values that could drive the observed variability in the outputs. Figure 7 shows the inferred parameter values which are used in the model predictions in Figure 5. Since we assume all cells share the same mechanistic model underlying the hERG currents our inferred cell-specific model parameters capture the cell-to-cell variability, or rather experiment-to-experiment variability. In Figure 7, our inferred parameters are plotted against manual patch parameters (shown as orange dots/red squares), measured at a slightly lower (room) temperature, from Beattie et al. (9); our identified parameters are broadly in alignment with manual patch results. This agreement gives us further confidence that our high-throughput method is reproducible and biophysically meaningful. We can also see that there is more variability in some parameters than others, also seen in the previous study (9).

**Figure 7:**
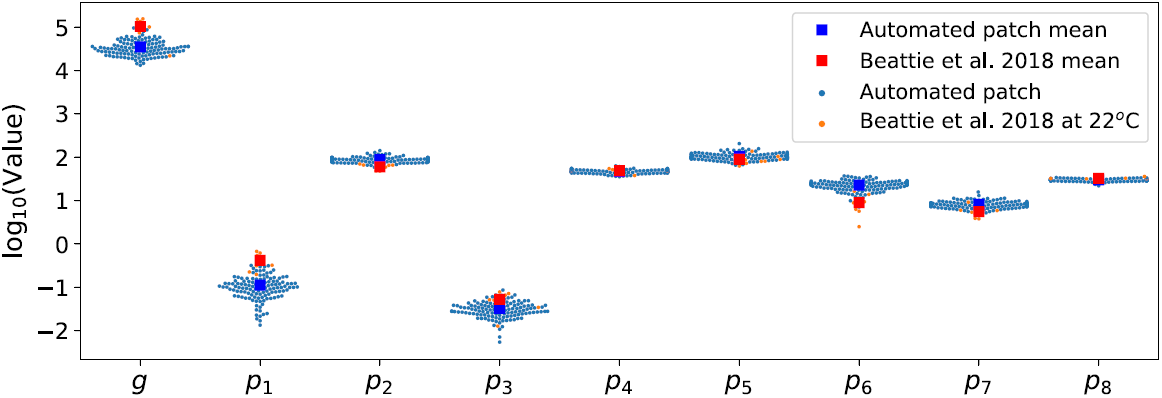
Cell-specific model parameters at around 25 °C. The inferred parameter values shown here are obtained from the staircase protocol calibration and are also the parameters used in the model predictions in Figure 5. It also shows the manual patch obtained parameters (orange), measured at around 22 °C, from Beattie et al. (9). The inferred kinetic parameter values from the automated high-throughput system are broadly consistent with the manual patch measurements.

### A hierarchical Bayesian modelling approach to characterise well-to-well variability

We applied the hierarchical Bayesian model to analyse the variability within the experimental recordings and correlations between inferred well-to-well parameter sets. The result of applying our hierarchical model is shown in Figure 8. The measurement uncertainty for the parameters of *each individual well* is shown with a marginal posterior distribution — the coloured histograms. Most of the parameters give a narrow credible interval, which reinforces our certainty in the information content of the calibration protocol. Many of the marginal posterior distributions of the individual wells overlap — that is, we cannot distinguish between the two sets of parameters given our uncertainty in them. However, some of the individual marginal posterior distributions are distinct from each other, demonstrating considerable variability *between wells*.

**Figure 8:**
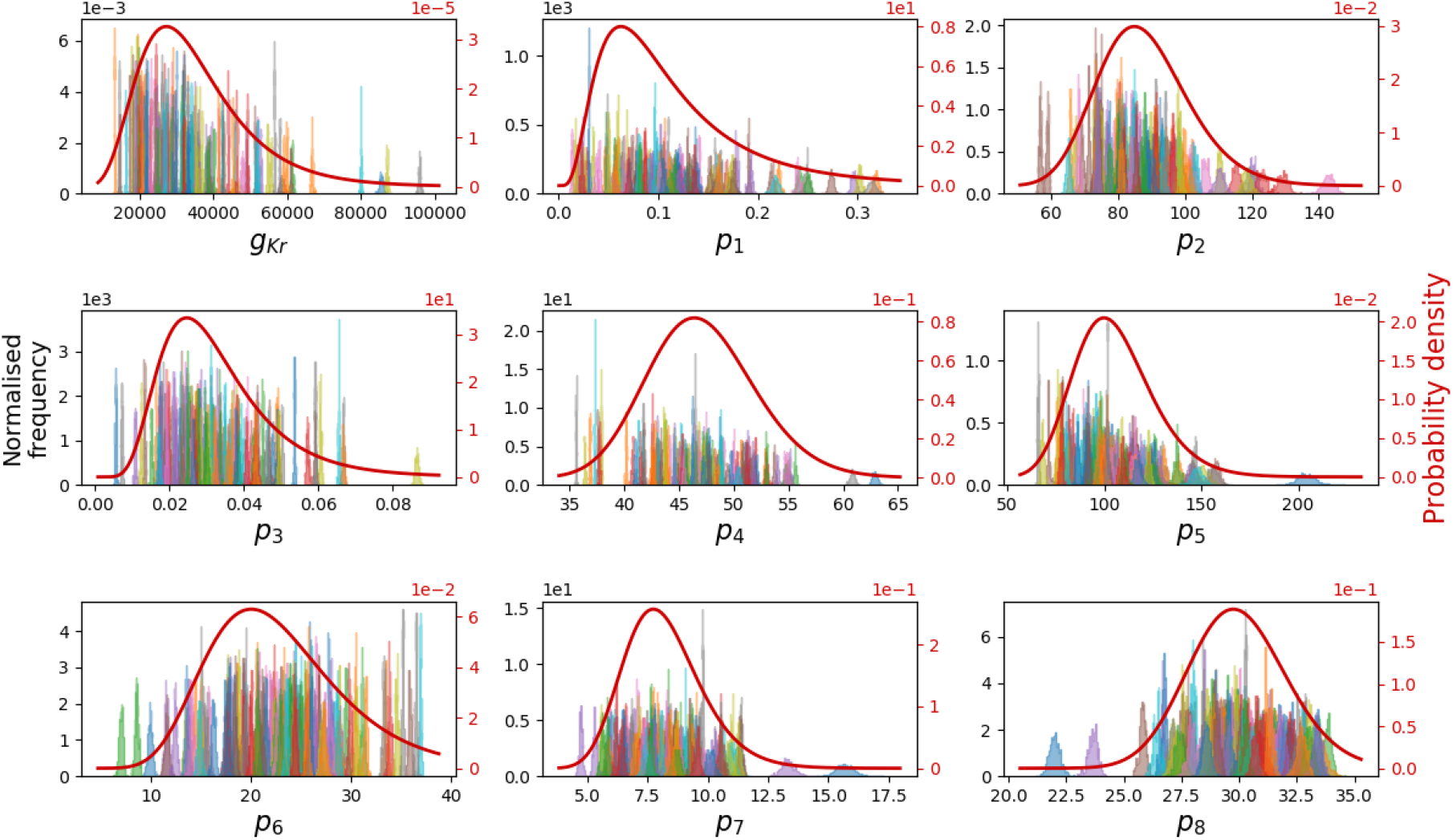
The marginal distributions from the hierarchical Bayesian model for all model parameters. **Left y-axis:** Shows the normalised marginal histograms (probability densities) for each parameter, representing the 124 individual experiments. All of them shows a narrow distribution which implies good confidence in our inferred parameters for the individual experiment. **Right y-axis:** The marginal posterior predictive distributions *p*(***θ*** | …), which are assumed to follow a multivariate log-normal distribution for each parameter, are shown as red curves. They are the inferred underlying distribution between cells for each of the parameters.

The power of the hierarchical Bayesian model can then be used to summarise and capture the experiment-to-experiment variability. The hyperparameters of the model describe both the mean ***µ*** and (co-)variance ****Σ**** of parameter sets across wells, with experimental uncertainty taken into account. We estimated this posterior predictive distribution (Eq. 9) from the samples of hyperparameters, and its marginal distributions are shown for each parameter as the red curves in Figure 8. This distribution can be used to *predict* the likelihood and variability of parameter sets from further wells in future experiments.

Finally, we utilised the hierarchical Bayesian model to investigate the correlation *between model parameters across different wells*. In the sampled hyperparameters, the covariance matrix ****Σ**** reveals any correlation between our model parameters. The typical assumption concerning the variability of parameters is that parameters are independent, i.e. in the covariance matrix all entries except the diagonal are zero. In the upper triangle (orange) of Figure 9, we compare our inferred correlation between parameters (calculated using Eq. 8) with this common assumption (black vertical dashed lines). It is obvious that there are many entries where zero is outside our credible interval, which is equivalent to showing that the independence assumption is not supported by our findings.

**Figure 9:**
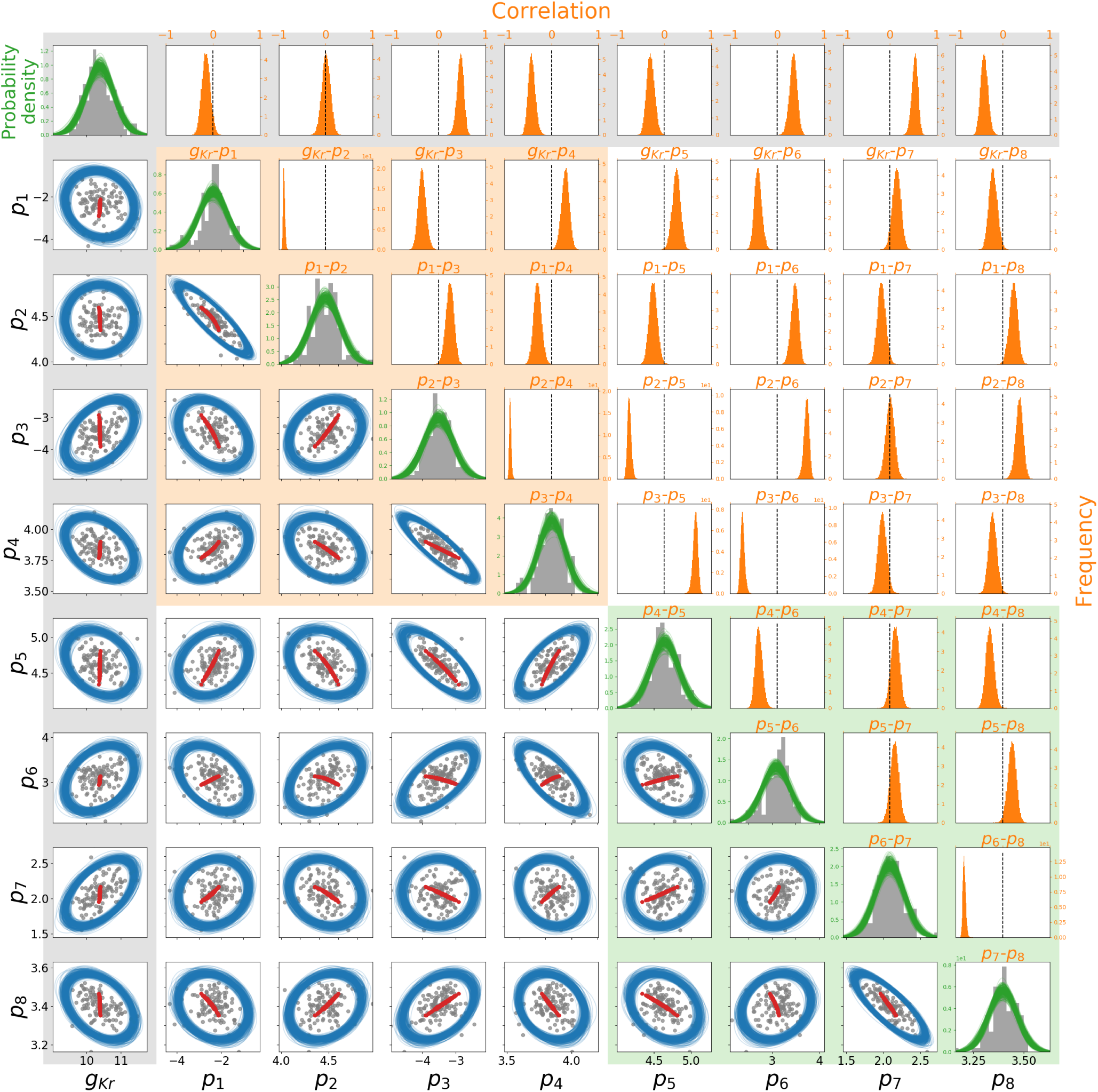
The inferred correlation in model parameters across experimental wells. All parameter values shown here are in the natural log-scale. The posterior mean parameters of each of the 124 individual cells are shown in grey (dots and histograms). **Lower triangle (blue):** The 95% credible region boundary for each pair of parameters. Each ellipse is reconstructed from one sample of the hyperparameters from the MCMC chain of size 10^5^, for clarity only 200 credible region samples are shown here. Simulated voltage error offset (described in the Discussion) is shown as red dots. **Diagonal (green):** the sampled posterior probably density functions before integration to give *p*(***θ*** | …), shown in detail in Figure 8. **Upper triangle (orange):** The marginal histograms for each entry of the correlation matrix defined by Eq. 8. The common assumption of independence (correlation of zero) is shown as black vertical lines for comparison. The shadings in the background indicate how these parameters relate to the model structure: the orange box contains the gates *a* in model, green box contains gate *r*, and grey relates to the conductance.

To visualise the correlation between parameters better, the 95% credible regions for each pair of parameters are shown in the lower triangle (blue) of Figure 9; plotted against the scatter plot of the 124 cells individual posterior mean parameters (shown on a log-scale). Each blue ellipse is reconstructed from a sample of hyperparameters, where the contour of the 95% credible region of the two-variate marginal distribution defined by the hyperparameter sample is shown; capturing most of our individual posterior mean parameters appropriately. In this plot, a perfect circle implies there is no correlation between the pair of parameters. However, we can clearly see that most of our pairwise parameters show an elliptical shape, which means some degree of correlation between the pairwise parameters exists. This strongly suggests that correlations between parameters are embedded in the experiment-to-experiment variability. We further discuss explanations for such observed correlations in the Discussion section.

## DISCUSSION

In this study, we have developed a short, high information-content staircase voltage clamp protocol for *I*_Kr_ that is applicable in automated high-throughput patch clamp systems, and used a mathematical model to characterise channel kinetics by fitting its parameters to recordings made under this new protocol. This study will advance future ion channel model development and model selection, and forms a basis for improved screening of ion channel kinetics, under different conditions, mutations or pharmaceutical compounds.

Here, we no longer use I-V or τ-V relations to characterise hERG kinetics, but rather we use a mechanistic model and its parameterisation to capture our knowledge of channel kinetics. An optimised voltage protocol, which is short and has a high information-content, was used to calibrate/parameterise the hERG kinetics model. The benefits of this approach are threefold. First, it is much easier to obtain a stable measurement, and one can worry less about current “rundown” during the protocol as compared to traditional I-V and τ-V protocols. Second, given its short duration, it is easy to repeat the measurement, to examine within-cell reproducibility/variability. Third, our staircase protocol can be used to rapidly create *cell-specific models of kinetics* (which is much harder to do using the more time-consuming traditional I-V and τ-V protocols).

We have shown that our 15 second staircase protocol can be performed in an automated high-throughput system. We have found that each of the resulting 124 models is consistent with previous manual patch clamp results (limited to 9 cells) (9), implying that these methods are reproducible. We can now easily produce large data sets for further analysis, which is usually difficult, if not impossible, to achieve with manual patch clamp. The predictions of the cell-specific models are not perfect, as we examined in Figure 6, and there may be room for improvement in terms of the model structure and further optimisation of the calibration protocol. But we are able to calibrate our model to the extent that it can replicate both experimental training data *and* predict validation data very well (Figure 4). Our models can predict the current response to the physiologically-relevant action potential protocols, demonstrating that our *I*_Kr_ models could be useful in predicting cardiac electrical activity in both healthy and arrhythmic situations (9). This provides assurance that our *cell-specific models*, which are constructed in a *high-throughput* manner, have great potential for future uses.

For example, our method can potentially be adapted and used to investigate not only how much the hERG channel is blocked by a drug, but also how that drug influences channel kinetics. This might be useful for the CiPA initiative, as both automated high-throughput systems and *in silico* modelling constitute the core of the initiative (24, 25). Our approach may give us a better understanding of the pharmacological properties of drugs in the screening process and hence a better pharmaceutical safety assessment. We can also incorporate the cell-to-cell or experiment-to-experiment variability in the *in silico* modelling as part of the uncertainty quantification for safety-critical predictions (14). Furthermore, such rapid characterisation using high-throughput systems can benefit precision and personalised medicine. For example, when using human induced pluripotent stem cell-derived cardiomyocytes (hiPSC-CMs) in personalised medicine, as described in (26), characterisation of ion current kinetics may need to be taken into account, in order to tailor accurate cell line-specific models.

With our 124 *cell-specific* hERG models, we are able to study experiment-to-experiment variability in the hERG channel. Such experiment-to-experiment variability is captured using our hierarchical Bayesian model, where the posterior predictive distribution is constructed and describes the underlying variability of the parameters (Figure 8). Instead of using a series of current-voltage (I-V) and time constant-voltage curves, here we evaluate the variability of the observed hERG channel kinetics using mathematical model parameters. The variability in the parameter values predicts the observed variability of the channel kinetics. In addition, we can use our posterior predictive distribution to predict what might happen in future experiments, based on the observed experiments.

### The source of variability

We have successfully quantified the variability *between wells* via our inferred model parameters. However, the underlying cause of this variability is an open question. There are possibilities at two extremes. One is that the variability is truly cell to cell, and ion channel kinetics do vary because of different intracellular conditions, one may speculate in terms of gene expression, the presence of subunits, phosphorylation states, or suchlike. The other possibility is that ion channel kinetics are precisely identical in each cell, but there are some experimental artefacts, varying between wells, that are causing the observed variability in parameters from each well. Below we discuss hints in our results as to which of these extremes is the leading cause of variability.

As mentioned in the rationale of the staircase protocol, the 100 ms ramp at 14.41 s was introduced to estimate experimentally the hERG reversal potential 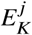 in each of the *j* wells, for details see Figure S2. Figure 10 shows an example of our 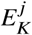 estimation using the ramp and a histogram of 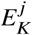 values estimated from the 124 wells.

**Figure 10:**
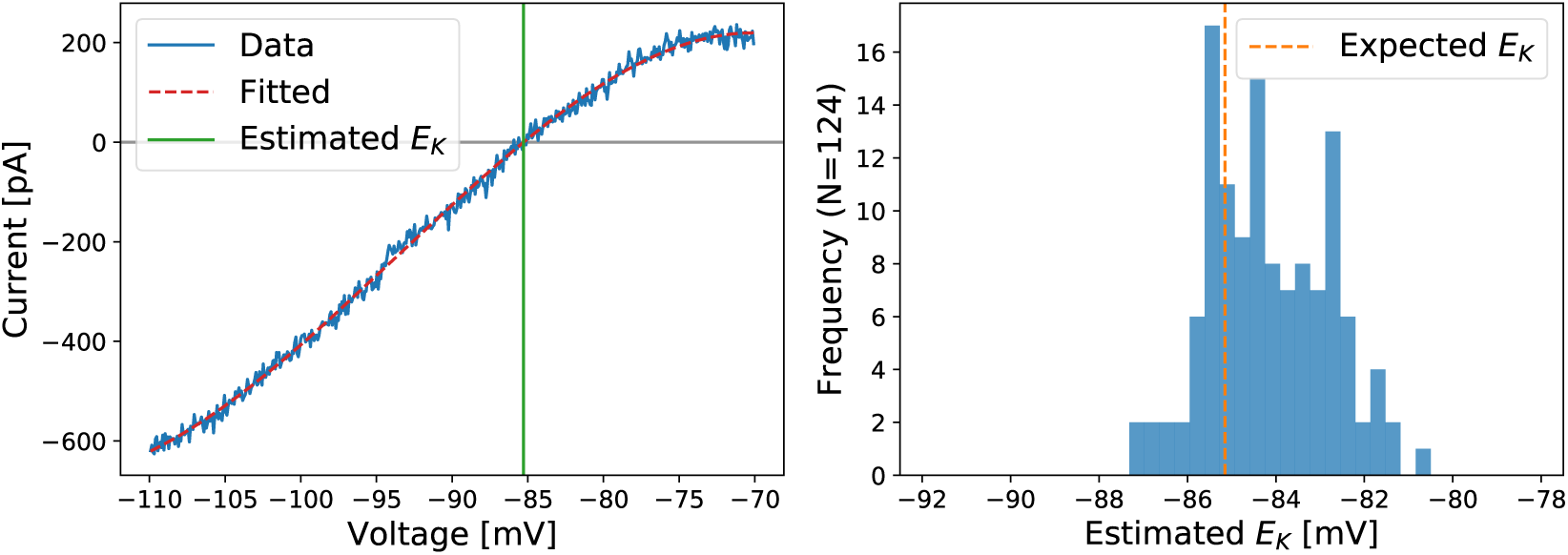
**Left:** An example of the current-voltage relationship plotted for the last ramp in the staircase protocol, and how it is used to estimate the *E*_*K*_ reversal potential value for *I*_Kr_in one well. **Right:** Histogram of *E*_*K*_ values estimated using the ramp technique. The *E*_*K*_ values here were estimated from the same 124 wells used in the main results. The dashed orange vertical line shows the expected *E*_*K*_ calculated directly from temperature and concentrations using the Nernst equation (Eq. 2).

Our obtained histogram of 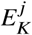 values is distributed close to our theoretical *E*_*K*_ from the Nernst equation (Eq. 2), with a standard deviation of 1.36 mV. Since all of our measurements were performed on one 384 well plate, they shared the same extra- and intra-cellular solutions and were recorded at (almost) the same temperature. We would therefore expect the real variability in reversal potential to be much smaller than this observed variability.

A hypothesis then, is that reversal potential *E*_*K*_ really occurs at the Nernst calculated value, and observed deviations from this inferred from the ramp provide an estimate for a ‘voltage error’ in the applied voltage clamp: 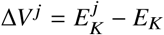, perhaps due to an imperfect compensations of other external effects, such as junction potentials and electrode offsets. We can investigate this hypothesis via the model: by applying a staircase protocol with voltage error offsets of Δ*V* ^*j*^ estimated from each of the 124 cells, generating synthetic data from these voltage clamps, and then re-fitting parameters.

Figure 9 (Lower triangle) shows the results of our voltage error offset simulations in red dots. If there was an error in the applied voltage clamp in each well, then we would expect to see parameters appearing to co-vary along the red lines (made up of individual dots/fits) in Figure 9. The observed primary parameter covariance directions and magnitudes from this procedure (red lines) align suspiciously well with much of the observed variability in the experiments (blue ellipses inferred from grey dots).

This is strong circumstantial evidence — a smoking gun — suggesting that the majority of the observed variability in parameters may be due to well-well variability in patch-clamp artefacts rather than cell-cell variability in ion channel kinetics. Building a more complete mathematical model of such patch clamp artefacts is part of our future plans. We should also note that despite patch clamp artefacts being an apparent cause of parameter variability, they are not necessarily larger artefacts in this automated system than might be expected in manual patch clamp.

Finally, if we were to believe that the observed variability here arises from experimental artefacts, then only the uncertainty in the top-level mean parameter vector ***µ*** in the hierarchical Bayesian model is representative of our uncertainty in the underlying physiology. That is, the variability of top-level mean parameter vector ***µ*** should be included in future *physiological* studies, for example in the second part of this study Lei et al. (27), or when embedding an ion channel model within an action potential model, whilst the full posterior predictive distribution should be used only when predicting the results of future patch clamp *experiments*.

## CONCLUSION

In this study, we have demonstrated the feasibility and practicality of using a 15-second staircase protocol to study and characterise hERG channel kinetics on an automated high-throughput system. We calibrated the hERG model to our staircase protocol for 124 hERG cells. Our 124 cell-specific variants of the hERG model are able to predict 8 other protocols with a high accuracy including physiologically-inspired action potential-like voltage clamps. Using a hierarchical Bayesian modelling approach, we provide a quantitative description of the variability and uncertainty within our 124 cell-specific models.

With our rapid characterisation techniques and the hierarchical Bayesian modelling approach, we have opened a new gateway to study parameter correlations between cells and investigate experimental variability. We have found that some model parameters are strongly cross-correlated, but not all. This result may hint at the origin of the variability and requires further investigation. In future, we aim to design protocols to allow high-throughput systems to be used to investigate not only how much the hERG channel is blocked by a drug, but also the kinetics of drug binding and whether the drug influences underlying channel kinetics.

## AUTHOR CONTRIBUTIONS

CLL, MC, DJG, LP, GRM & KW designed the research. CLL, LP & KW carried out the experiments. CLL, MC, DJG & GRM designed the computational analysis. CLL wrote simulation codes, performed the analysis, and generated the results figures. All authors wrote and approved the final version of the manuscript.

## ACKNOWLEDGEMENTS

We thank Nanion for assistance with exporting current trace data.

This work was supported by the Wellcome Trust [grant numbers 101222/Z/13/Z and 212203/Z/18/Z]; the Engineering and Physical Sciences Research Council and the Medical Research Council [grant number EP/L016044/1]; and the Biotechnology and Biological Sciences Research Council [grant number BB/P010008/1]. CLL acknowledges support from the Clarendon Scholarship Fund; and the EPSRC, MRC and F.Hoffman-LaRoche Ltd for studentship support via the Oxford Systems Approaches to Biomedical Science Centre for Doctoral Training. MC and DJG acknowledge support from a BBSRC project grant. GRM acknowledges support from the Wellcome Trust & Royal Society via a Sir Henry Dale Fellowship and a Wellcome Trust Senior Research Fellowship.

LP and KW are employees of F.Hoffman-LaRoche Ltd. and KW is a shareholder.

## SUPPORTING MATERIAL

An online supplement accompanies this article. All codes and data are freely available at https://github.com/CardiacModelling/hERGRapidCharacterisation.

