## Supporting Material for "Rapid characterisation of hERG channel kinetics I: using an automated high-throughput system"

### Supporting Material for:
### Rapid characterisation of hERG channel kinetics I

**Chon Lok Lei[1], Michael Clerx[1], David J. Gavaghan[1], Liudmila Polonchuk[2], Gary R. Mirams[3], and Ken Wang[2]**

[1]Computational Biology, Department of Computer Science, University of Oxford, Oxford, United Kingdom
[2]Pharma Research and Early Development, Innovation Center Basel, F. Hoffmann-La Roche Ltd., Basel, Switzerland
[3]Centre for Mathematical Medicine and Biology, School of Mathematical Sciences, University of Nottingham, Nottingham, United Kingdom

#### Contents

**S1 Details of voltage clamp protocol** **1**

**S2 Ramps in the staircase protocol** **2**

**S3 Electrophysiology solutions** **4**

**S4 Recording techniques** **5**

**S5 Automated quality control** **5**

**S6 Synthetic data studies** **8**

**S7 Sweeps comparison** **17**

**S8 Posterior predictive quantification** **18**

**References** **21**

#### S1 Details of voltage clamp protocol

##### S1.1 Calibration: Staircase protocol

The full protocol is comprised of a 250 ms step at holding potential of −80 mV, followed by a 50 ms 'leak step' at −120 mV, and a 400 ms 'leak ramp' from −120 mV to −80 mV, before a 200 ms back at holding potential. This was followed by a 1 s 'activation step' at 40 mV and a 500 ms 'closing step' at −120 mV, before returning to holding potential for 1 s. Then the 9.5 s staircase portion of the protocol (the details is described below), before a return to holding potential for 500 ms. Finally, it was followed by a reversal potential estimation portion which is composed of a 500 ms step to 40 mV, and a 10 ms step to −70 mV to remove capacitance effect, then followed by a 100 ms 'reversal potential ramp' starting from −70 mV to −110 mV, before a 390 ms step to −120 mV, and return to holding potential for 500 ms.

The staircase portion of the protocol consists of a range of 500 ms steps up and down as discussed in main text. It is comprised of two sets of steps, the first set alternates between $V_{step,1}$ and $V_{step,2}$, each for 500 ms. There are 5 different $V_{step,1}$ and $V_{step,2}$; $V_{step,1}$ ranged from −40 mV to 40 mV, and $V_{step,2}$ ranged from −60 mV to 20 mV, both in 20 mV increments. The second set alternates between $V_{step,3}$ and $V_{step,4}$, each for 500 ms. There are 5 different $V_{step,3}$ and $V_{step,4}$; $V_{step,3}$ ranged from 40 mV to −40 mV, and $V_{step,4}$ ranged from 0 mV to −80 mV, both in 20 mV decrements.

This protocol is shown in Figure S1A. A time series version of the full protocol is available at https://github.com/chonlei/hERGRapidCharacterisation/blob/master/protocol-time-series/protocol-staircaseramp.csv.

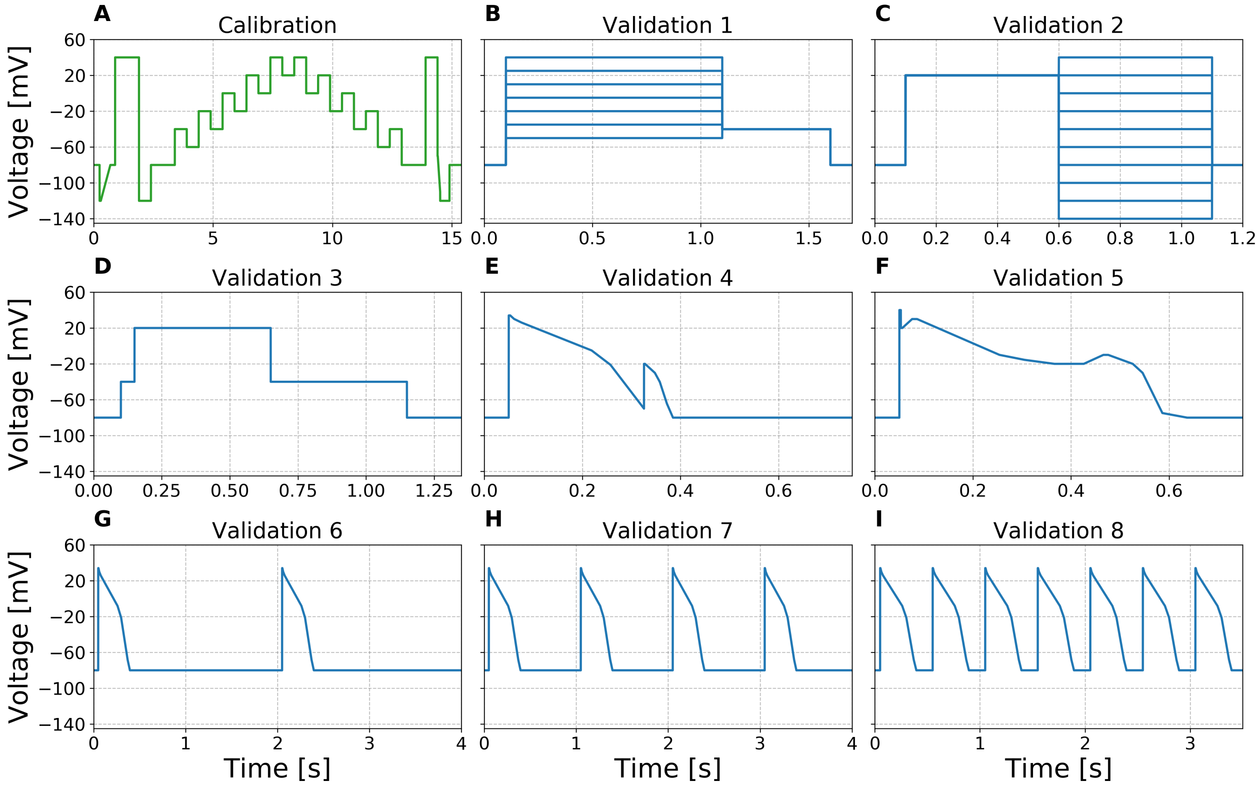


**Figure S1.** All voltage clamp protocols used in the study, from **A** to **I** are (green) our newly developed staircase protocol, (blue) the activation current-voltage (I-V) protocol, the steady-state inactivation I-V protocol, the hERG screening protocol, the early afterdepolarization (EAD)-like protocol, the delayed afterdepolarization (DAD)-like protocol, and the action potential (AP)-like protocol with beating frequency 0.5 Hz, 1 Hz and 2 Hz. All protocols are shown with the same voltage axes for comparison; however due to different time scale, each of them has its own time axis.

#### S1.2 Validation 1: Activation I-V protocol

From the initial period 100 ms at holding potential of $-80$ mV, a step to $V_{step}$ for 1 s, followed by a 500 ms step to $-40$ mV, before a 100 ms step back to holding potential; this was repeated 7 times with a different $V_{step}$ on each repeat. $V_{step}$ ranged from $-50$ mV to 40 mV in 15 mV increments. This protocol is shown in Figure S1B.

#### S1.3 Validation 2: Steady-state inactivation I-V protocol

From the initial period 100 ms at holding potential of $-80$ mV, a step to 20 mV for 500 ms, followed by a step to $V_{step}$ for 500 ms, before a 100 ms step back to holding potential; this was repeated 10 times with a different $V_{step}$ on each repeat. $V_{step}$ ranged from $-140$ mV to 40 mV in 20 mV increments. This protocol is depicted in Figure S1C.

#### S1.4 Validation 3: hERG screening protocol

From the initial period 100 ms at holding potential of $-80$ mV, a step to $-40$ mV for 50 ms, and a step to 20 mV for 500 ms, followed by a step to $-40$ mV for 500 ms, before a 200 ms step back to holding potential. This protocol is shown in Figure S1D.

#### S1.5 Validation 4-8: EAD-like, DAD-like, APs-like protocols

Details are described in Table S1, and each protocol is shown in Figure S1E-I respectively.

***EAD-like protocol***

| Type | V [mV] | Duration [ms] |
|---|---|---|
| Step | -80 | 50 |
| Step | 34 | 3 |
| Ramp | 30 | 8 |
| Ramp | 26 | 15.2 |
| Ramp | -5 | 142.6 |
| Ramp | -21 | 38.4 |
| Ramp | -70 | 68.6 |
| Step | -20 | 2 |
| Ramp | -30 | 20 |
| Ramp | -40 | 10 |
| Ramp | -65 | 15 |
| Ramp | -80 | 12 |
| Step | -80 | 15.2 |
| Step | -80 | 350 |

***DAD-like protocol***

| Type | V [mV] | Duration [ms] |
|---|---|---|
| Step | -80 | 50 |
| Step | 40 | 3 |
| Step | 20 | 3 |
| Ramp | 30 | 20 |
| Step | 30 | 10 |
| Ramp | -10 | 168 |
| Ramp | -15.5 | 50.6 |
| Ramp | -20 | 61.2 |
| Step | -20 | 60 |
| Ramp | -10 | 40 |
| Step | -10 | 10 |
| Ramp | -20 | 50 |
| Ramp | -30 | 20 |
| Ramp | -75 | 40.5 |
| Ramp | -80 | 50 |
| Step | -80 | 13.7 |
| Step | -80 | 100 |

***Single AP-like protocol***

| Type | V [mV] | Duration [ms] |
|---|---|---|
| Step | -80 | 50 |
| Step | 34 | 3 |
| Ramp | 30 | 8 |
| Ramp | 26 | 15.2 |
| Ramp | -8 | 183.6 |
| Ramp | -21 | 39 |
| Ramp | -68 | 70.5 |
| Ramp | -80 | 25.2 |
| Step | -80 | — |

**Table S1.** Details of the EAD-like (validation 4), DAD-like (validation 5), APs-like (validation 6-8) protocols. It shows as a sequence of steps and ramps that approximates different types of action potential shapes, as these are the only available settings in the automated machine used. The voltage (V) in the type Ramp represents the final targeted voltage that the ramp finishes, starting from the previous voltage within the given duration; for example, the first ramp in the EAD-like protocol means it starts from 34 mV and ramps to 30 mV in 8 ms. The single AP-like protocol shows the protocol for one unit AP-like protocol that repeats in 0.5 Hz, 1 Hz, and 2 Hz.

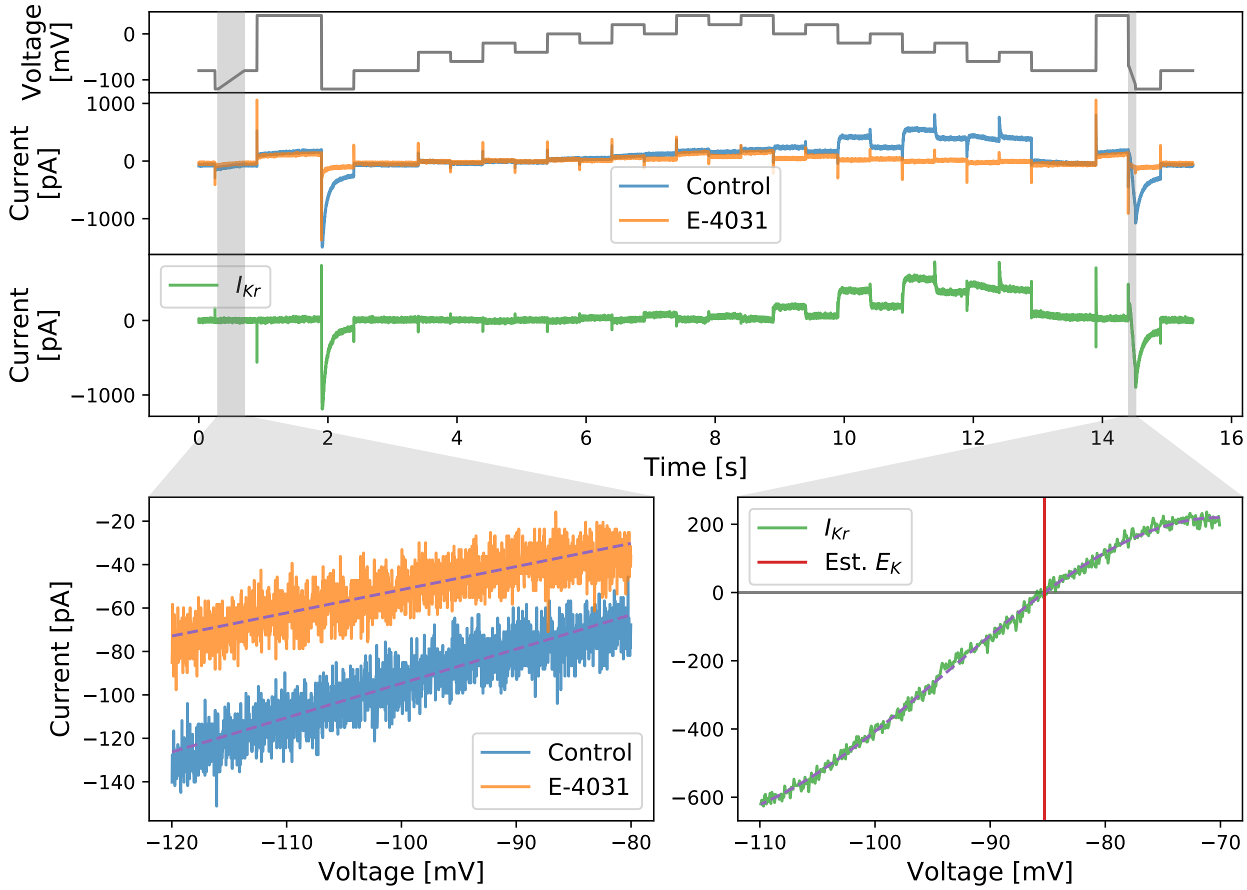


**Figure S2.** Two ramps (greyed out sections) implemented in the staircase protocols. The first ramp is designed to estimate the leak current; the second ramp is designed to experimentally estimate the $E_K$ value. Top three panels show the staircase voltage clamp protocol (grey), the raw currents before (blue) and after (orange) E-4031 application, and the estimated $I_{\mathrm{Kr}}$ (green) traces, respectively. Below shows the I-V curves measured during the two ramps. Linear regressions were applied to each of the I-V relation in the first ramp, shown as dashed lines. Third order Polynomial regression was applied to the I-V curve in the second ramp, shown as dashed lines.

#### S2 Ramps in the staircase protocol

As discussed in the main text, protocol design, the two ramps implemented in the staircase protocol are designed to estimate the leak current and to experimentally estimate the $E_K$ value. Figure S2 shows an example of using the two ramps to estimate the leak current and the $E_K$ value. The top three panels show the staircase voltage clamp protocol (grey), an example of raw currents before (blue) and after (orange) E-4031 application, and the corresponding estimated $I_{\mathrm{Kr}}$ (green) traces, respectively. The greyed out sections highlight the two ramps in the staircase protocol. Bottom left shows the I-V curves of the two raw currents measured under the first ramp. Linear regressions were applied, and the results are shown as dashed lines, where the fitted slope and y-interception point were used to estimate the leak current parameters (Eq. 12 in the main text). Bottom right shows the I-V curve of the leak-corrected, E-4031 subtracted $I_{\mathrm{Kr}}$ measured under the second ramp. A third order polynomial regression was applied, and the result is shown as dashed line. The $E_K$ value was then estimated as the x-interception point, shown as red vertical line.

#### S3 Electrophysiology solutions

The compositions of all the electrophysiology solutions, including both the external solutions (bath solutions) and the internal solution (equivalent to the pipette solution in manual patch clamp), are shown in Table S2. External solutions were added in the following order: first 'fill chip' solution to the measurement chip, and the suspended hERG cells, then the 'seal enhancer' solution for enhancing the seal by forming CaF crystal around the cells (note they have extra high concentration of $Ca^+$, so we need to reduce/dilute it later), followed by adding the extracellular 'reference' solution for $Ca^+$ dilution. All the voltage clamp measurements were performed after adding all these external solutions.

The solutions were added sequentially to the wells, by removing half of the previous solutions from the wells each time. Therefore, the final ratios of the external (extracellular) solution are 1:1:2 — proportions of 0.25 of the 'Fill Chip' concentrations, 0.25 of the 'Seal Enhancer' concentrations, and 0.5 of the 'Reference' concentrations, as shown in the 'Final Extracellular' solution in Table S2.

| **Solution** | | *Intracellular* | *Fill Chip* | *Seal Enhancer* | *Reference* | *Final Extracellular* |
|---|---|---|---|---|---|---|
| **pH value (titrated with)** | | *pH 7.2 (KOH)* | *pH 7.4 (NaOH)* | *pH 7.4 (HCl)* | *pH 7.4 (HCl)* | |
| **Osmolarity [mOsm]** | | *260-300* | *300-330* | *290-330* | *290-330* | |
| **Chemicals** | **Source / Cat#** | **[ ] in mM** | **[ ] in mM** | **[ ] in mM** | **[ ] in mM** | **[ ] in mM** |
| NaCl | Merck / K38447104807 | 10 | 150 | 80 | 80 | 97.5 |
| KCl | Merck / K36782536 | 10 | 4 | 4 | 4 | 4 |
| KF | Acros Organics / 201352500 | 100 | — | — | — | — |
| $MgCl_2$ | Merck / A914133908 | — | 1 | 1 | 1 | 1 |
| $CaCl_2$ | Acros Organics/ 349615000 | — | 1.2 | 5 | 1 | 2.05 |
| HEPES | Applichem A1069 | 10 | 10 | 10 | 10 | 10 |
| Glucose | Fluka / 49159 | — | 5 | 5 | 5 | 5 |
| NMDG | Fluka 66930 | — | — | 60 | 40 | 35 |
| EGTA | Fluka / 03778 | 20 | — | — | — | — |
| Sorbitol | Sigma / S1876 | — | — | — | 40 | 20 |

**Table S2.** Electrophysiology solutions for hERG assay on the Nanion SyncroPatch 384PE machine, all solutions are sterile filtered. All hERG cells were suspended in 1/3 Extracellular Fill Chip Solution + 2/3 Hanks' Balanced Salt Solution (HBSS).

#### S4 Recording techniques

All experiments were performed with Nanion SyncroPatch 384PE machine with software PatchControl384PE (v. 1.5.6 Build 22) and current traces data were exported using their complementary software DataControl384 (v. 1.5.0 Customer Release). Temperature was controlled by Nanion temperature control unit with software PE384TemperatureControl. The machine comes with a measurement chip consists of 364 wells, with 16 rows by 24 columns.

#### S5 Automated quality control

Here we present a more detailed selection results of our quality control which does not require any manual intervention. The full details of our automated quality control criteria are summarised in Table 1 in the main text. A well must pass all the listed criteria in order to be selected.

In Figure S3, we break down the selection results and show the results of each criterion in our automated quality control. On the left, the bar chart shows the number of well removed by each quality control criterion. In which there were 22 'no cell' wells, where the system decided there was no valid estimation of $R_{seal}$, $C_m$, and $R_{series}$ and did not process recording. Our three **QC1** criteria are used as part of the automated high-throughput machine quality control, which can eliminate up to 46 wells out of the 201 well that we manually decided to remove. We then added the other criteria to improve the selection process, which allow us to eliminate a total of 173 wells, and achieved a positive predictive value of >86 %. On the right, we show the

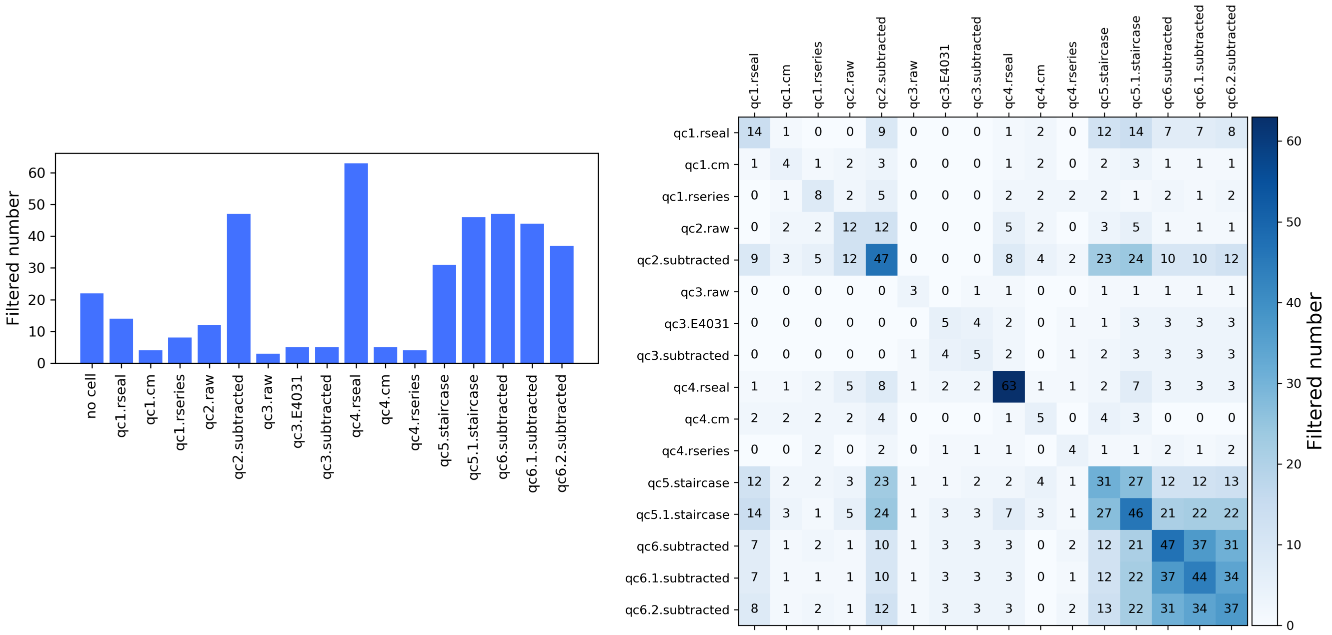


**Figure S3.** Selection results of each criterion from our automated quality control. **(Left.)** Showing the number of wells filtered out by each quality control criterion as bar chart. **(Right.)** Showing the number of wells filtered out by both the row and column criteria. The automated high-throughput machine also has some simple quality control implemented, which are our three **QC1** criteria.

number of wells commonly removed by any pair of criteria. This shows that most of our criteria are quite independent, and are assessing different features of the recordings.

We note that our automated quality control can achieve a positive predictive value of $>86\,\%$, here we show 6 typical examples of the ones that we manually removed. Figure S4 shows a comparison of the good recordings and our manually removed bad recordings. Top panel shows our staircase protocol. Then we show 3 good recordings (green) and 6 manually removed bad recordings (orange/red). We found our manually removed recordings fall into two main categories, as coloured, orange and red.

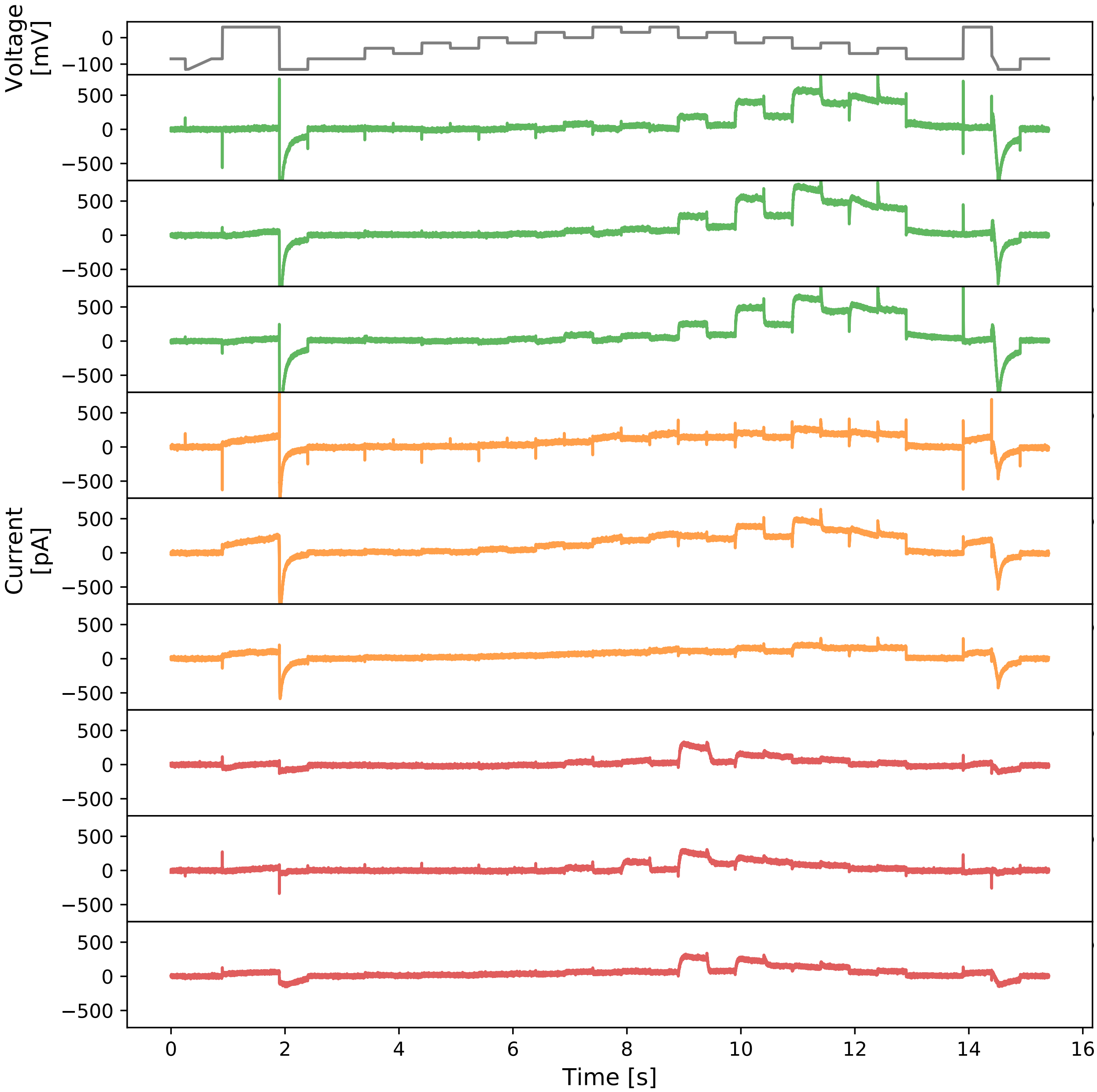


**Figure S4.** A comparison of the good recordings and our manually removed bad recordings. Top panel shows our staircase protocol. Following are 3 good recordings (green) and 6 manually removed bad recordings (orange/red). We found our manually removed recordings fall into two main categories, as coloured, orange and red.

### S6 Synthetic data studies

#### S6.1 Introduction

We perform a synthetic data study prior to implementing the actual experiments for two reasons: design of protocols and test of analysis.

First, given the mathematical model of the hERG channel, we are able to deduce what would be the best protocols to tease out the kinetics of the underlying model. This can alleviate a common issue, identifiability when undertaking model fitting. Because typically one constructs the problem into inverse problems when trying to parameterise mathematical models, however the identifiability issue arises because of the poorly informed experimental data. Therefore we utilise a synthetic study to design and optimise protocols to have sufficient information for rapid characterisation.

Second, we are able to test our analysis technique, to ascertain whether it is robust enough for our purpose — to recover the parameters of the model given the data. Most of our ion channel models can be written as

$$I = f(V, t; \theta, I_0), \tag{S1}$$

where $I$ is the current (output of the model, the observable in experiments), $V$ is the voltage, and $\theta$ is the vector of parameters within the model. The models are usually formulated as differential equations which therefore requires initial conditions $I_0$. The dependency on initial conditions $I_0$ can usually be eliminated by running the model long enough to reach a (pseudo-)steady state. Then with our analysis techniques, given the output $I$ with inputs $V$ and $t$, we aim to infer the values of the parameters $\theta$, hence the overall process is termed an inverse problem. Therefore, we generate synthetic data (with added synthetic noise) with some 'true' parameters $\theta^{\text{true}}$, and we ask, how confident are we in our inferred parameters?

#### S6.2 Methods

##### *S6.2.1 Generating synthetic data*

We generate synthetic data by simulating the current $I$, with some fixed known parameter sets $\{\theta^{\text{true}}\}$, voltage protocol $V_{prt}(t)$, initial values $I_0$, and sampling time (time-step) $\Delta t$.

First, the choice of $\{\theta^{\text{true}}\}$ could be arbitrary, but we used the parameters identified from a previous study[1] (Table F11 Cell #5), $\theta^{\text{lit}}$, to utilise prior knowledge. We generated $\{\theta^{\text{true}}\} = \{\theta^{\text{true},1}, \theta^{\text{true},2}, \cdots, \theta^{\text{true},N_e}\}$ with each $\theta^{\text{true},j} = (\theta_1^{\text{true},j}, \theta_2^{\text{true},j}, \cdots, \theta_N^{\text{true},j})^T$ sampled from

$$\theta_i^{\text{true},j} \sim \mathcal{N}(\theta_i^{\text{lit}}, \rho_i^2), \tag{S2}$$

where $i = 1, 2, \cdots, N$ for $N$ parameters in the model, and $(\cdots)^T$ represents the transpose. $\mathcal{N}$ denotes the normal distribution and $\rho^2$ is the variance for which we chose a value of $\rho_i = 0.2|\theta_i^{\text{lit}}|$. That is, we assume that we performed $N_e$ experiments (recordings), and there exists variability between experiments. Assuming each experiment was performed identically, then the variability that we are simulating is *cell-to-cell variability*.

We can take the notion of variability further, by removing the assumption of independence *between* model parameters. We assume there exists an underlying correlation between each model parameter, which can be described by a covariance matrix $\Sigma$. Therefore we can rewrite the underlying distribution of the parameters as being taken from a (covarying) multivariate normal distribution, that is

$$\theta^{\text{true},j} \sim \mathcal{N}(\theta^{\text{lit}}, \Sigma). \tag{S3}$$

The correlation between parameters using the correlation matrix is then defined as

$$\text{corr}(\theta) = \text{diag}(\Sigma)^{-1/2}\ \Sigma\ \text{diag}(\Sigma)^{-1/2}, \tag{S4}$$

where $\text{diag}(\cdot)$ denotes the matrix of the diagonal entries and its $(i,i)$ entry is chosen to be $\rho_i^2$. We randomly generated the correlation matrix that satisfies the positive semi-definite condition for this synthetic data study.

Second, we fix the voltage $V$ of the model at $V_{prt}(t)$, which is the staircase protocol that we developed for the high-throughput systems. Third, for the initial values $I_0$, we ran the model at $V = -80\,\text{mV}$ for a long period (100 s), to allow the model to settle at its steady state at $V = -80\,\text{mV}$. Since we are able to mimic this in the actual experiments, we assume the model does not depend on the choice of $I_0$, that is $I \approx f(V,t;\theta)$.

Finally, we add synthetic noise which follows a normal distribution with a mean of zero and standard deviation $\sigma$ (i.e. $\sim \mathcal{N}(0,\sigma^2)$) to the simulated traces with $\Delta t = 0.5\,\text{ms}$. We chose $\sigma$ at a reasonable scale, $\sigma = 11\,\text{pA}$, to mimic the high frequency noise observed from the experiments.

##### S6.2.2 Inferring parameters

To infer the parameters, we use a two-step approach. Firstly, we use a global optimisation algorithm, CMA-ES[2], to identify the parameters. Secondly, we run Markov-chain Monte Carlo (MCMC) to explore and quantify the uncertainty of the identified parameters.

In the CMA-ES optimisation, we used the sum of squares error measure of the whole trace as our objective function. To alleviate any potential issues arising due to a constrained objective function, we applied a transformation $g$ that maps the positively constrained model parameters $\{\theta_i\}$, with $\theta_i \in [0,\infty]$, to $\{\phi_i\} \in \mathbb{R}^N$, an unconstrained search space for optimisation, which is simply a log-scale transformation:

$$\theta_i = g^{-1}(\phi_i) = e^{\phi_i}. \tag{S5}$$

We then further considered the physical constraints for the rate constants in the kinetics parameters[1], which has the form $k = A\exp(BV)$. For parameters of the form $A$, $[\theta_i^{\min}, \theta_i^{\max}]$ is chosen to be $[10^{-7}, 10^3]\,\text{ms}^{-1}$; and for parameters of the form $B$, $[\theta_i^{\min}, \theta_i^{\max}]$ is chosen to be $[10^{-7}, 0.4]\,\text{mV}^{-1}$.

For the MCMC, we used a population MCMC[3] algorithm with adaptive Metropolis[4] algorithm as the base sampler. The starting point of the population MCMC was chosen to be the CMA-ES inferred parameters. As a good practice, the population MCMC was repeated 3 times to ensure the convergence of the MCMC chains. We chose the posterior measure to be

$$p(\phi,\sigma|\mathbf{y}) = \frac{p(\phi)p(\mathbf{y}|\phi,\sigma)}{p(\mathbf{y})} \propto p(\phi)p(\mathbf{y}|\phi,\sigma), \tag{S6}$$

$$p(\phi) \sim \mathscr{U}(\phi^{\min},\phi^{\max}), \tag{S7}$$

$$p(\mathbf{y}|\phi,\sigma) = \frac{1}{\sqrt{2\pi\sigma^2}} \exp\left(-\sum_k \frac{\left(f(V_{prt},t_k;g^{-1}(\phi)) - \mathbf{y}|_{t_k}\right)^2}{2\sigma^2}\right). \tag{S8}$$

Here, $\mathbf{y}$ is the data and $\mathbf{y}|_{t_k}$ denotes the data at time $t_k$. The likelihood, $p(x|\phi)$, in Eq. S8 is the Gaussian noise version of the sum of square difference measure used in the CMA-ES.

##### S6.2.3 Hierarchical Bayesian model

In order to infer the correlation *between* model parameters, $\text{corr}(\theta)$ in Eq. S4, the mean, and the variability between cells, we used a multi-level modelling technique which works under the Bayesian framework, known as a hierarchical Bayesian model. This allows us to combine all the results from each individually performed experiment to inform the prediction of future experiments.

A schematic of our hierarchical Bayesian model structure is shown in Figure S5. The full hierarchical Bayesian model is

$$\begin{aligned}\mathscr{L}\left(\mu,\Sigma,\left\{\phi_j,\sigma_j\right\}_{j=1}^{N_e} | \left\{\mathbf{y}_j\right\}_{j=1}^{N_e}\right) \propto \prod_{j=1}^{N_e} p\left(\mathbf{y}_j|\phi_j,\sigma_j\right) \\ \times p\left(\left\{\phi_j,\sigma_j\right\}_{j=1}^{N_e} |\mu,\Sigma\right) \\ \times p(\mu,\Sigma) \times \prod_{j=1}^{N_e} p(\sigma_j),\end{aligned} \tag{S9}$$

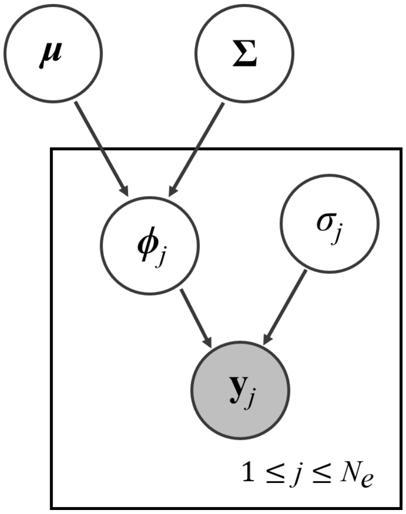


**Figure S5.** Hierarchical Bayesian model showing parameter dependency for combining multiple experiments. $\mu, \Sigma$ are the hyperparameters of the hierarchical model which represent the mean and covariance matrix, respectively, of the individual 'low-level' parameters, $\{\theta_j, \sigma_j\}_{j=1}^{N_e}$ are the set of individual 'low-level' parameters for each of the $N_e$ measurements in the high-throughput experimental recordings $\{\mathbf{y}_j\}_{j=1}^{N_e}$. The parameters in the box repeat for multiple wells and are indexed as the $j^{\text{th}}$ experiment (or dataset). All parameters, and their probability distributions, are inferred from the shaded variable $\mathbf{y}_j$, the experimental data. Prior distributions are required for the parameters with no inward-pointing arrows.

where all symbols have their usual meaning as defined above, $\mathscr{L}$ is the full posterior, and $\mu, \Sigma$ are the hyperparameters of the hierarchical model which are the means and covariance matrix of the model parameters. We assume the model parameters follow a multivariate log-normal distribution, thus the hyperparameters define the mean and covariance matrix of this distribution. The three terms in Eq. S9 are: 1. the likelihood of *all* the individual (low-level) experiments; 2. the likelihood of the hyperparameters; and 3. the priors of the hyperparameters (also known as 'hyper-priors') and the prior of $\sigma_j$ which we do not infer its hyperparameters.

For computational ease, we chose the prior of the hyperparameters to be a multivariate normal distribution for the $\mu$ and an inverse-Wishart distribution $\mathscr{W}^{-1}$ for the $\Sigma$, which is the respective conjugate prior. Suppose $N_e$ individual parameters $\{\theta_j\}_{j=1}^{N_e}$ have been observed, then we have

$$\left\{\ln \theta_j\right\}_{j=1}^{N_e} \sim \mathscr{N}(\mu, \Sigma), \tag{S10}$$

and with the conjugate prior

$$p(\mu, \Sigma) = p(\mu|\Sigma)p(\Sigma), \tag{S11}$$

where

$$p(\mu|\Sigma) \sim \mathscr{N}\left(\mu_0, \frac{1}{m}\Sigma\right), \quad \text{and} \quad p(\Sigma) \sim \mathscr{W}^{-1}(\Psi, \nu). \tag{S12}$$

$\mu_0, m, \Psi, \nu$ are the prior parameters, where $m, \nu$ are respectively the strength of the prior mean $\mu_0$ and $\Psi$ which determines the prior of the covariance $\Sigma$. Then the posterior distribution of the hyperparameters becomes

$$p\left(\mu \,|\, \Sigma, \left\{\ln \theta_j\right\}_{j=1}^{N_e}\right) \sim \mathscr{N}\left(\frac{N_e\bar{\theta} + m\mu_0}{n+m}, \frac{1}{m+N_e}\Sigma\right), \text{ and} \tag{S13}$$

$$p\left(\Sigma \,|\, \left\{\ln \theta_j\right\}_{j=1}^{N_e}\right) \sim \mathscr{W}^{-1}\left(\Psi + N_e S + \frac{N_e m}{N_e + m}(\bar{\theta} - \mu_0)(\bar{\theta} - \mu_0)^T, N_e + \nu\right), \tag{S14}$$

where

$$\bar{\theta} = \frac{1}{N_e}\sum_{j=1}^{N_e} \ln\theta_j, \quad \text{and} \tag{S15}$$

$$S = \frac{1}{N_e}\sum_{j=1}^{N_e} (\bar{\theta} - \ln\theta_j)(\bar{\theta} - \ln\theta_j)^T. \tag{S16}$$

We use the Metropolis within Gibbs (MwG)[5] sampling method to explore the full hierarchical Bayesian model. The number of parameters we have in Eq. S9 is $N(N+1)/2 + (N_e+1)N + N_e$. For our choice of hERG model and the size of the dataset, we are expecting $N = 9$ and $N_e > 100$. This gives us more than 1000 parameters for which we wish to infer probability distributions. It is computationally expensive and infeasible to use other standard algorithms, such as the population MCMC, and even MwG can be very time consuming. We therefore further simplify the MwG to approximate the full posterior sampling, which we have termed '*pseudo-MwG*'. We confirm that the pseudo-MwG can approximate the MwG very well in the results below.

Under our pseudo-MwG, we assume that the likelihoods of our individual experiments are unlikely to be affected by the top-level distribution, due to our information-rich staircase protocol having thousands of data points rather than the $\sim 100$ wells. We therefore separate the sampling steps between the likelihood of all the individual experiments and the likelihood of the hyperparameters. That is, we first *independently* sample the likelihood of each individual experiment, using population MCMC alogrithm. Then we sample the hyperparameters using Eq. S10–S16, where $\{\ln\theta_j\}_{j=1}^{N_e}$ become $\{\ln\theta_{j,l}\}_{j=1}^{N_e}$ which are the independently obtained $l^{\text{th}}$ samples of the individual experiments $j$. Note that this is only valid when the individual experiments are far more information-rich than the number of repeats. We check these assumptions in the results below.

To obtain the posterior predictive distribution $p(\theta|\cdots)$ which allows us to make prediction about how the future experiments would behave, where $(\cdots)$ indicates all other variables appear in Eq. S9, we use

$$p(\theta|\cdots) = \int_\Theta p(\theta|\Theta)p(\Theta|\cdots)\,\mathrm{d}\theta, \tag{S17}$$

where $\Theta = (\mu,\Sigma)^T$. This can be approximated by summing over the probability density functions which are defined by the samples of $\Theta$.

##### S6.3 Results/Discussion

###### S6.3.1 Single synthetic experiment

We start by showing the staircase protocol is information-rich enough to identify the 'true' parameter set in a synthetic data study using our protocol. Figure S6 shows the results of inferring model parameters on a synthetic experiment, where $\theta^{\text{true}} = \theta^{\text{lit}}$ obtained from a previous study[1] (Table F11 Cell #5). It shows the three independently sampled marginal posterior distributions of each parameter (first and third columns), with indications of the 'true' parameters $\theta^{\text{true}}$ (black dashed lines) which we used to generate the synthetic data, and the CMA-ES inferred parameters (red lines). Both the traces (second and fourth columns) and the three independently run posterior distributions show a good indication of the convergence of the MCMC chains. We are able to recover the 'true' parameters $\theta^{\text{true}}$ with high accuracy and a narrow credible interval using our inference techniques together with our developed staircase protocol. Therefore we are confident that, with both the high information-content protocol and the inference techniques, it is theoretically possible to infer all parameters of the model.

###### S6.3.2 Hierarchical synthetic experiments

Figure S7 shows the results of the synthetic data study using hierarchical Bayesian model with $N_e = 120$. It shows the marginal histograms of the model parameters for each individual experiment (left y-axis) and the marginal posterior predictive distribution (right y-axis, red lines). This synthetic data study is equivalent to have $N_e$ repeats of the same experiment. Unlike the single experiment study above, the implications of the obtained posterior predictive distribution $p(\theta|\cdots)$ are much more powerful and can be viewed in two ways.

First, we can see this as the underlying distribution that governs the parameters. That is, with this, we can try to understand – through the model – what the hERG channel is doing in the cells. To do so, we compare it with the 'true' underlying distribution

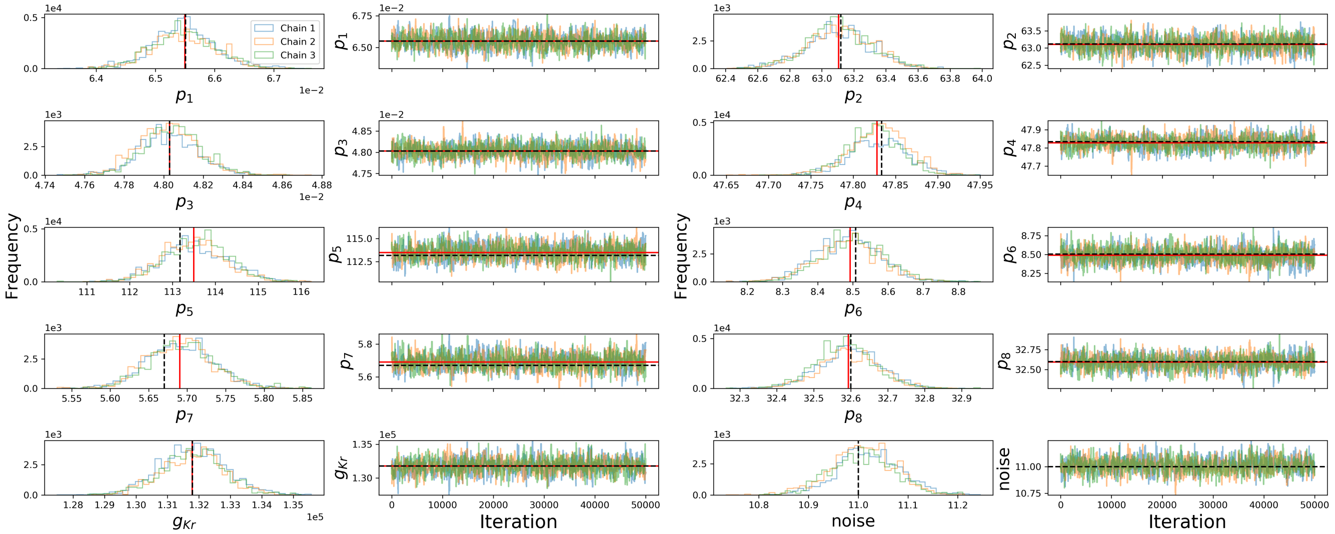


**Figure S6.** Parameter inference of single synthetic experiment, $N_e = 1$. **First, Third columns:** Show the marginal histograms of the posterior distribution of each parameter. **Second, Fourth columns:** The trace plots for our MCMC chains indicating that our MCMC chains have converged. Each panel shows the posterior distribution of 3 independently run MCMC, and their extremely good agreement assures the chains are well mixed. The true (synthetic) parameters are indicated as black dashed lines and the CMA-ES inferred parameters are shown as red lines.

of parameters (black dashed lines), i.e. the multivariate normal distribution in Eq. S3. The marginal posterior predictive distributions closely resemble the 'true' distribution. This indicates that we are able to recover the underlying distribution of the parameters with high accuracy too, and therefore we can rely on it to study the behaviour of hERG cells in actual experiments.

Second, as the name implies, this is a *predictive* distribution. That is, given the observed individual experiments, we infer a distribution which allows us to predict what might happen in a future experiment. To do this, we can view the posterior predictive distribution in Eq. S17 as $p(\theta_{N_e+1}|\cdots)$, where $\theta_{N_e+1}$ is our 'future' $(N_e+1)^{\text{th}}$ experiment that we perform. Therefore the distribution that we construct is able to tell us what is likely to happen in the future experiments — based on the observations from previous experiments.

We further investigate the correlation between parameters, by trying to recover the correlation matrix $\text{corr}(\theta)$ in Eq. S4. The posterior marginal histograms for each entry of the correlation matrix are shown in Figure S8 (upper triangle). The diagonal is by definition equal to 1, so they are not shown. All inferred marginal posterior distribution for each entry covers the true underlying correlation value (dashed black vertical lines). Therefore it shows us with confidence that our method is suitable for studying the relation *between* model parameters.

Figure S8 (lower triangle) shows the correlation between each pair of parameters. Each contour ring represents the 95% credible intervals of the joint distribution of the two parameters, for both the recovered (blue) and the true (dashed black) covariance matrices. As long as the main axis of the ellipse is not parallel to the x- or y-axis, it indicates the two parameters are not pairwise-independent. The diagonal shows the sampled predictive posterior distribution before integrated over to give $p(\theta|\cdots)$ shown in Figure S7. Again, it shows that we are able to recover the general shape of the underlying correlation with high accuracy.

In this synthetic study, the correlation matrix that we recovered may not make any physical sense – as we randomly generated it. However, in actual experiments, this correlation matrix tells us which parameters are intrinsically correlated. That is, if there exists any non-zero values, with a good credible interval, in the off-diagonal entries of the recovered correlation matrix, then this informs us how the parameters of model are related.

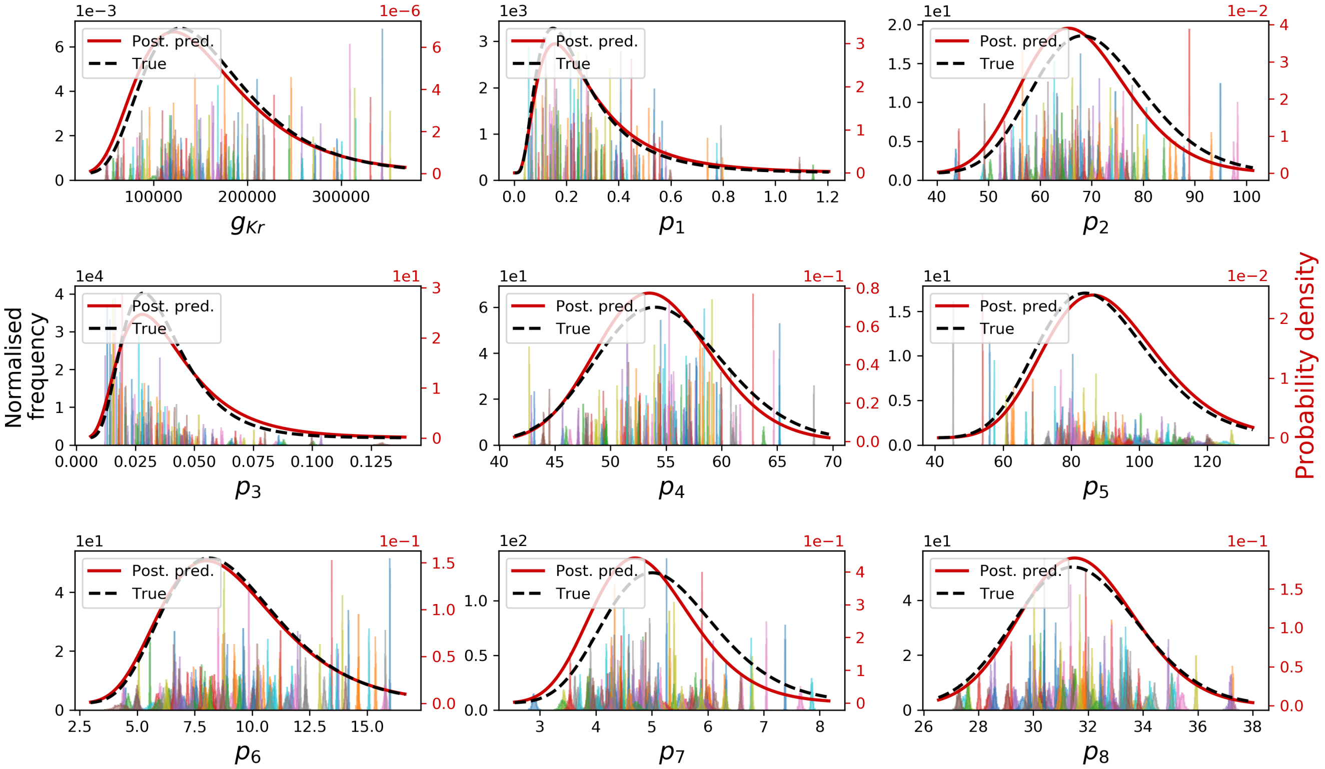


**Figure S7.** Parameter inference using the hierarchical Bayesian model on synthetic data, with $N_e = 120$. **Left y-axis**: the marginal histograms of the model parameters for each individual experiment. **Right y-axis**: the marginal posterior predictive distributions and the true probability density function that generates the parameters.

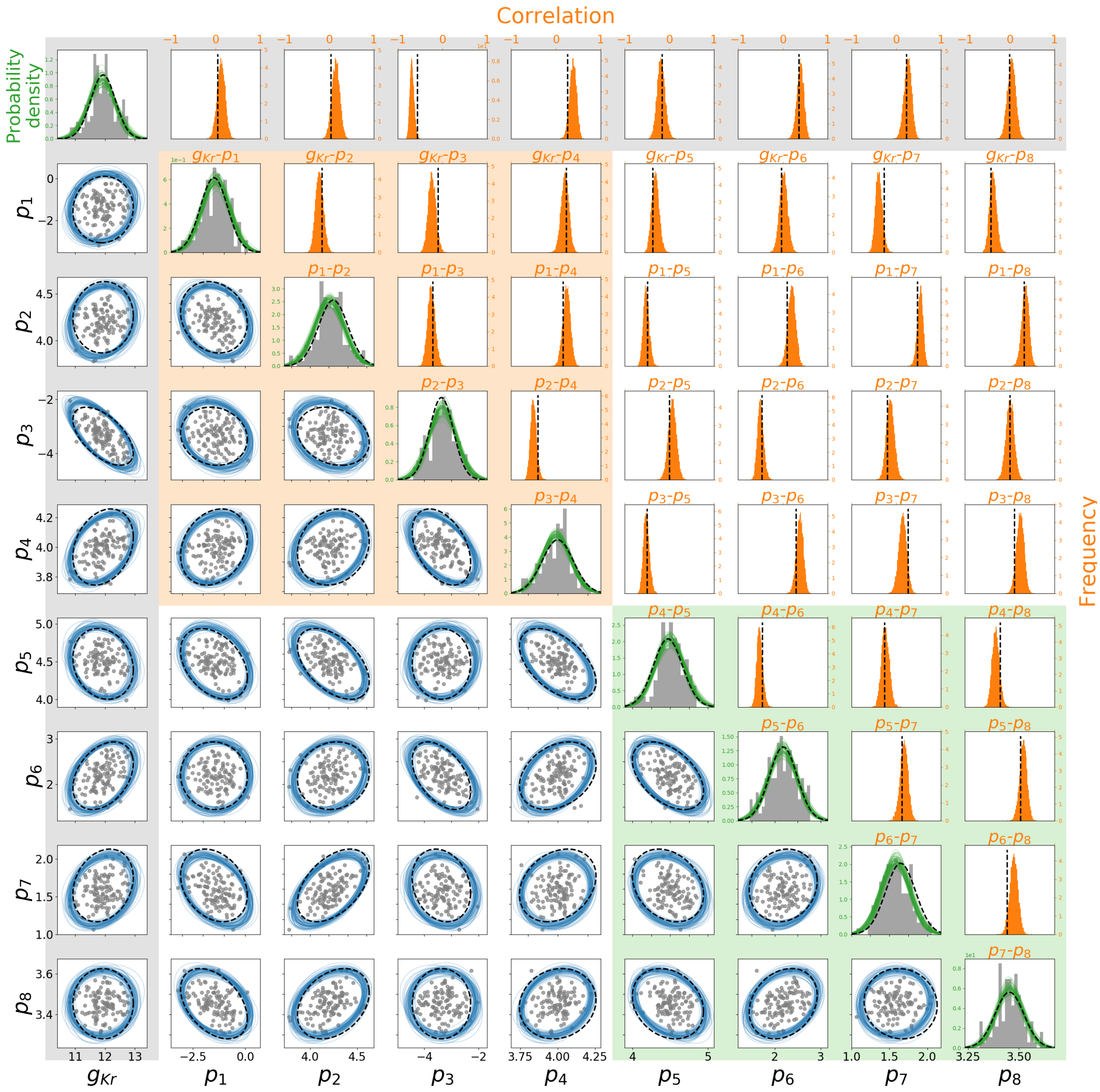


**Figure S8.** Parameter correlation inference using the hierarchical Bayesian model on synthetic data, with $N_e = 120$. **(Lower-triangle)** Showing the 95% credible intervals for each pair of parameters reconstructed from the sampled hyperparameters (blue) and the true distribution (red). **(Diagonal)** Shows the sampled predictive posterior probably density functions before integrated to give $p(\theta|\cdots)$ shown in Figure S7. The true 95% credible intervals and probably density function on the diagonal are shown in dashed black lines for comparison. Our synthetic data are shown as grey. **(Upper-triangle)** Shows the marginal histograms for each entry of the correlation matrix. The true correlation values are shown as dashed black vertical lines for comparison. The shadings in the background indicate how these parameters relate to the model structure: orange box belongs to the gates $a$ in model, green box gate $r$, and grey relates to the conductance.

**Comparing Pseudo-MwG to MwG** All the results above and those in the main text use pseudo-MwG method. Here we provide a brief comparison between our pseudo-MwG method and the MwG for approximating the posterior predictive distribution. We use $N_e = 30$ to demonstrate their similarity.

Figure S9 shows the posterior predictive distribution and the histograms of the individual experiments constructed from the pseudo-MwG method (solid lines/filled) and the MwG method (dashed lines/unfilled). It has the same style of plot as in Figure S7, where the left-axes show the marginal histograms and the right-axes show the marginal posterior predictive distributions. The posterior predictive distributions constructed from the pseudo-MwG and MwG look extremely similar. Therefore, with our staircase protocol as the likelihood of the low-level experiments, we are able to simplify our procedure to the pseudo-MwG without losing much accuracy comparing to the MwG algorithm.

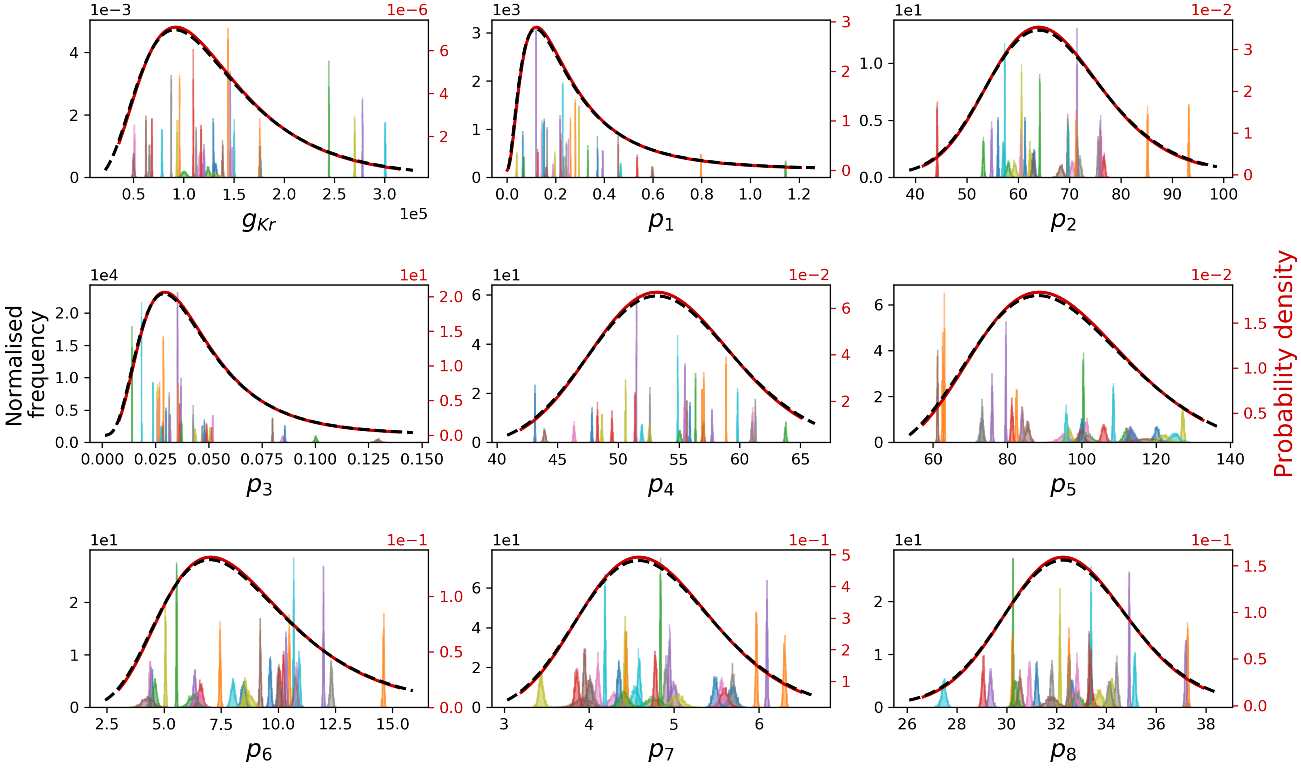


**Figure S9.** Comparing the hierarchical Bayesian model parameter inference on synthetic data using the pseudo-MwG (solid lines/filled) and the MwG (dashed lines/unfilled) methods, with $N_e = 30$. **Left y-axis**: the marginal histograms of the model parameters for each individual experiment. **Right y-axis**: the marginal posterior predictive distributions.

We also note that we can further simplify our pseudo-MwG, which we shall call it as *simplified pseudo-MwG*, given our information-rich staircase protocol. First we can see that the MCMC distributions, see e.g. Figure S9 marginal histograms, are really narrow relative to spread of each experiment parameters. By approximating these narrow distributions as single points (i.e. delta functions), we can then sample the hyperparameters using Eq. S10–S16, where $\{\ln \theta_j\}_{j=1}^{N_e}$ become point-estimates of the parameters of the individual experiments $j$. Figure S10 shows the posterior predictive distribution constructed from the *simplified* pseudo-MwG method (solid lines) and the MwG method (dashed lines). Again, the posterior predictive distributions constructed from the simplified pseudo-MwG and MwG look extremely similar. Therefore, we can further simplify our pseudo-MwG sampling scheme to estimate the full posterior-predictive distribution.

**Converging to the true distribution** We then check the performance of our method with different numbers of experiments/cells $N_e$, and confirm that the result converges to the correct answer. We calculate the score with root mean square error

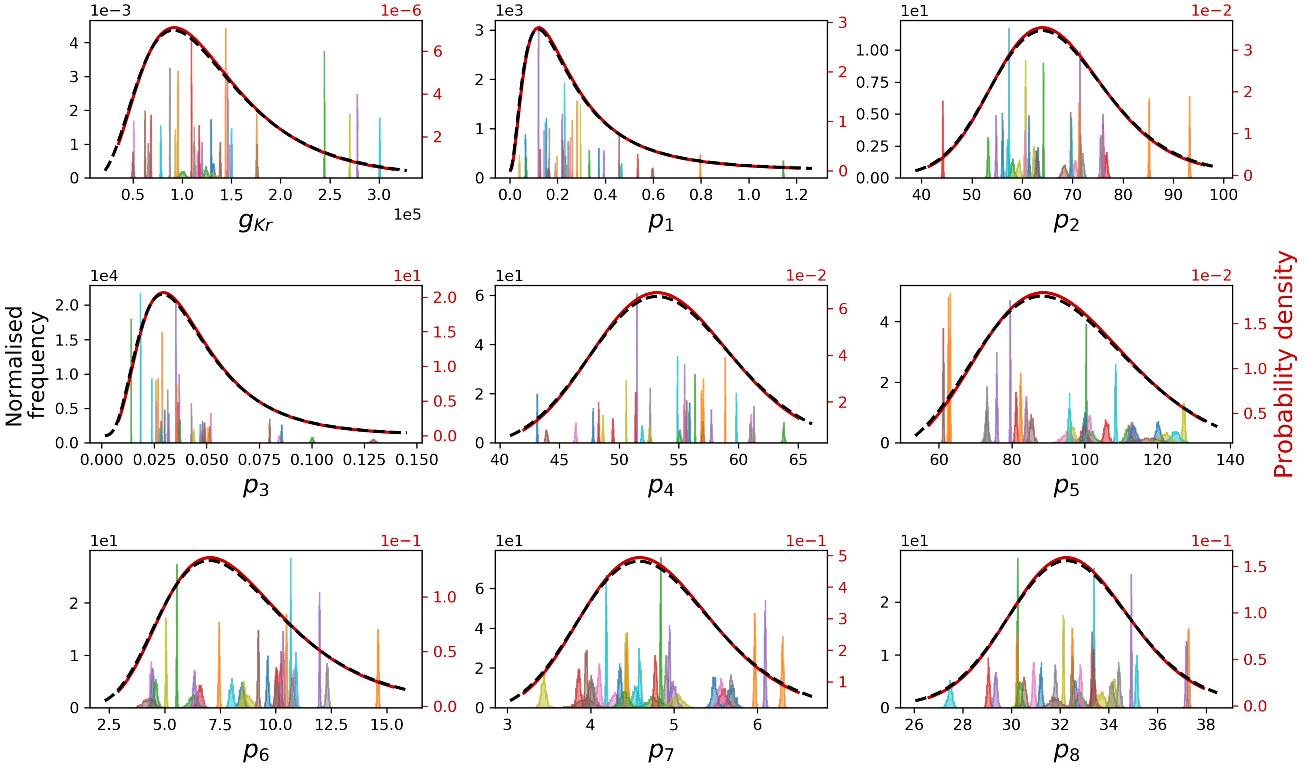


**Figure S10.** Comparing the hierarchical Bayesian model parameter inference on synthetic data using the *simplified* pseudo-MwG (solid lines) and the MwG (dashed lines) methods, with $N_e = 30$. **Left y-axis**: the marginal histograms of the model parameters for each individual experiment for the MwG method. **Right y-axis**: the marginal posterior predictive distributions.

(RMSE) for the correlation matrix, where

$$\text{RMSE of correlation} := \frac{1}{N}\sqrt{\sum_i^N \sum_j^N (\text{corr} - \text{corr}_{\text{true}})_{i,j}}, \tag{S18}$$

and its slight variant root mean square percentage error (RMSPE) for standard deviation, where

$$\text{RMSPE of std} := \sqrt{\frac{1}{N}\sum_i^N \frac{\text{std}^i - \text{std}^i_{\text{true}}}{\text{std}^i_{\text{true}}}}. \tag{S19}$$

We used RMSPE, instead of normal RMSE, for standard deviation to avoid different parameter magnitudes from dominating the calculation.

Figure S11 shows the RMSPE of the standard deviation (left) and RMSE of the correlation (right) as function of the numbers of experiments/cells $N_e$. For the RMSPE of the standard deviation, Figure S11 (Left), we repeated the above analysis with $N_e = 20, 30, \ldots, 120$ and 125. We can clearly see that the RMSPE of the standard deviation decreases as $N_e$ increases. Hence it is convincing that our method is converging to the true answer in the synthetic data studies.

For the RMSE of the correlation, Figure S11 (Right), we further test the convergence rate of the RMSE value. To run sufficiently large $N_e$, we simplified our procedure by running only the top-level of the hierarchical Bayesian model, i.e. the *simplified* pseudo-MwG as described above. With this, we ran $N_e$ up to $2 \times 10^4$. We plotted both axes in natural-log scale. We then applied a linear regression, in which a slope of $-0.516$ is obtained. Therefore, we conclude that convergence rate of the RMSE of the correlation is roughly consistent with $\propto 1/\sqrt{N_e}$. We also expect the likely errors in our experiments, with $N_e = 124$, is about 6.4 %, shown as grey lines.

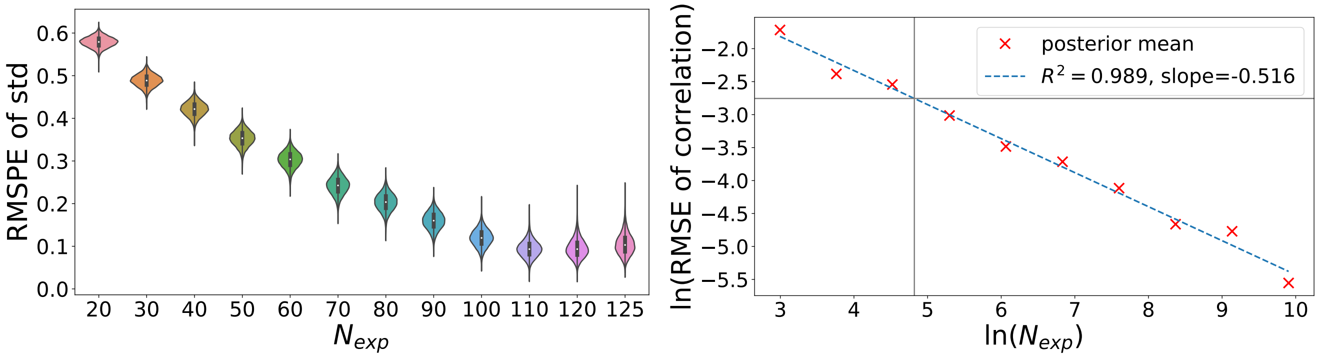


**Figure S11.** The RMSPE of standard deviation (left) and RMSE of correlation (right) as function of the numbers of experiments/cells $N_e$. Each violin plot and posterior mean is constructed using $10^4$ samples. Grey lines show where $N_e = 124$, with an RMSE value of 0.064.

#### S7 Sweeps comparison

Here, we check the reproducibility of our results *in the same cells*. We performed the same fitting procedure to the second sweep of our staircase protocol (calibration protocol) recording. First, to assess, if any, intrinsic (or intra-cell) variability[6] in our recordings; and second, to ensure our results are reproducible and biologically meaningful.

Figure S12 shows the fitted parameters comparison between the first sweep (sweep 1) and the second sweep (sweep 2) for all $N = 124$ cells. The line of identity is plotted as grey dashed lines. The two sets of parameters broadly agree, therefore it is convincing that our results are reproducible within the same cells. The intrinsic variability in our recordings are quite small, compared to the extrinsic or experiment-to-experiment variability. Therefore, our analyses focus on the observed experiment-to-experiment variability, and the intrinsic variability are assumed to be negligible.

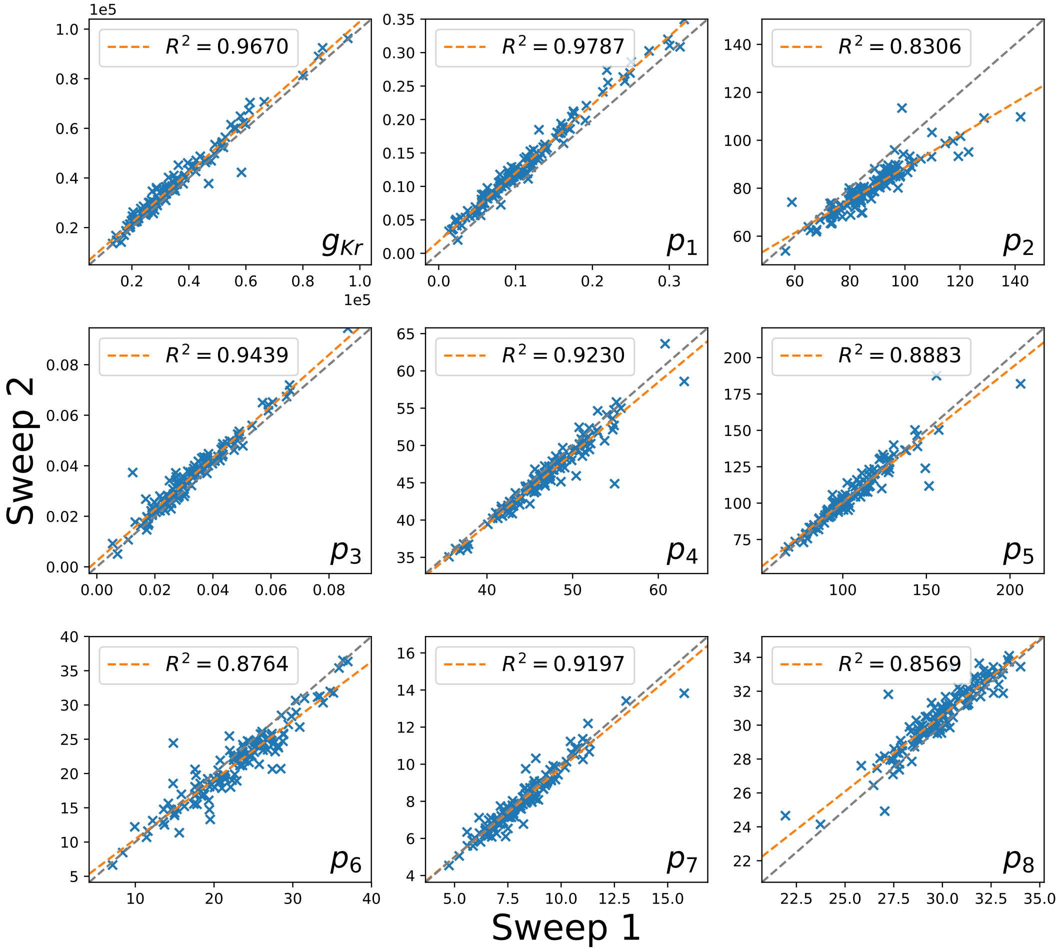


**Figure S12.** Comparison of the fitted parameters between the first sweep (sweep 1) and the second sweep (sweep 2). Grey dashed lines show the line of identity. The two sets of parameters broadly agree, therefore it is convincing that our results are reproducible within the same cells.

#### S8 Posterior predictive quantification

We quantify the goodness of the posterior predictive distribution from our hierarchical Bayesian model, compared to the 124 individual experiments, by means of a quantile-quantile (Q-Q) plot and a probability–probability (P-P) plot. The Q-Q (or P-P) plot is a graphical method for comparing two probability distributions, in our case the 124 individual experiments and our posterior predictive distribution $p(\theta|\cdots)$, by plotting their quantiles (or cumulative distributions) against each other.

Note that this is a good test of the LogNormal distribution because we used the pseudo-MwG method, and the individual level parameter fits were not allowed to shift to meet a LogNormal by design as a hierarchical model would generally behave.

Figure S13 and S14 show the Q-Q and P-P plots respectively. In both figures, for each parameter, the marginal posterior predictive distributions are plotted against the posterior mean of the 124 cells. We applied linear regression, shown as orange lines, and they all lie very close to the line of identity (grey dashed lines). These analyses support our results and suggest our posterior predictive distribution, defined by Eq. S17, is a very good description to the distribution of the data.

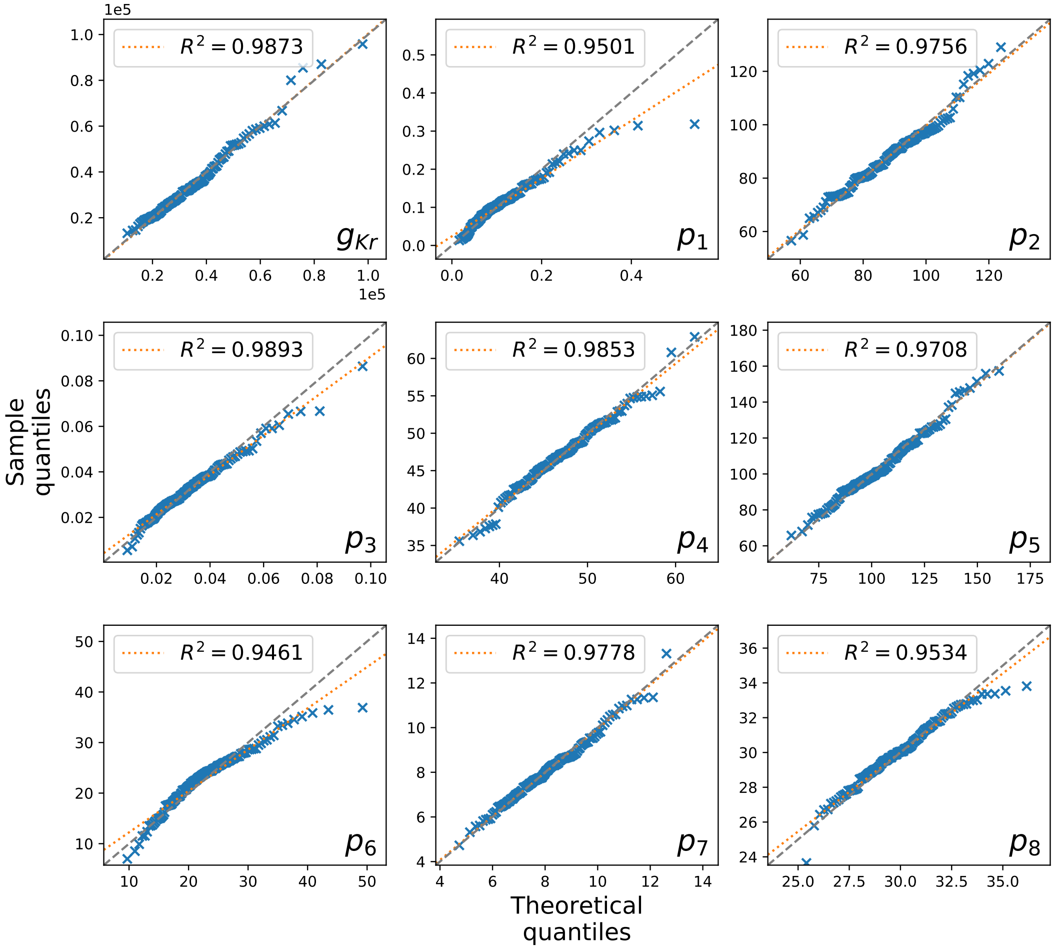


**Figure S13.** Quantile-quantile (Q-Q) plot of the 124 individual experiments and our posterior predictive distribution. For each parameter, the quantiles of the marginal posterior predictive distribution (theoretical quantiles) are plotted against the quantiles of the posterior mean of the 124 cells (sample quantiles).

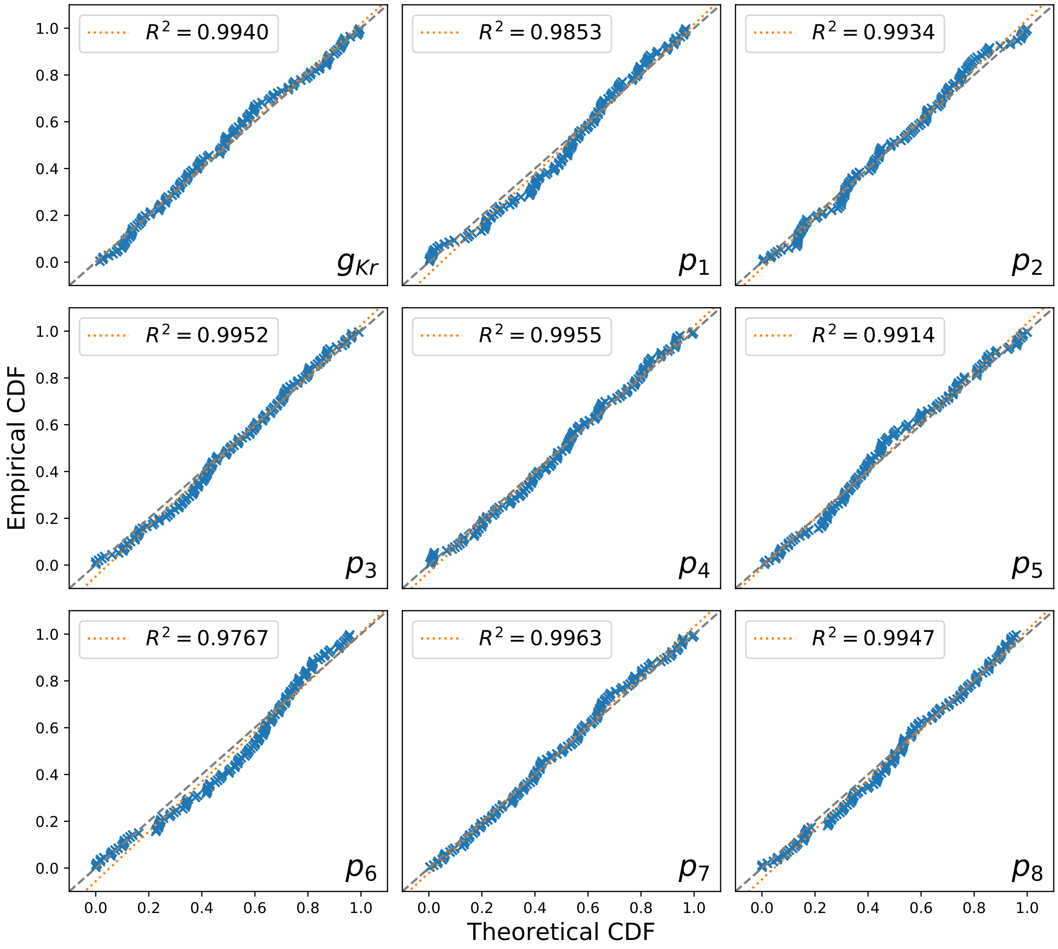


**Figure S14.** Probability–probability (P-P) plot of the 124 individual experiments and our posterior predictive distribution. For each parameter, the cumulative distribution of the marginal posterior predictive distribution (theoretical CDF) are plotted against the cumulative distribution of the posterior mean of the 124 cells (empirical CDF).
